# Harnessing the Intrinsic Dynamics of Biological Neural Networks for Reservoir Computing

**DOI:** 10.64898/2026.09.08.750274

**Authors:** Samitha Somathilaka, Jacob Clouse, Sasitharan Balasubramaniam

**Author notes:** Contributing authors. These authors contributed equally to this work (Jacob Clouse, Sasitharan Balasubramaniam).

## Abstract

Living neuronal networks exhibit nonlinear, recurrent, and evolving dynamics that make them promising substrates for reservoir computing, yet their computational use is complicated by spatial heterogeneity, spontaneous state transitions, and biological nonstationarity. Here, we investigate whether the native dynamics of a neuronal culture can be characterized and harnessed as a living reservoir without deliberately modifying the underlying recurrent network. Using multielectrode-array recordings and electrical perturbations, we characterize spontaneous population dynamics, validate channel-adaptive spike detection, and evaluate reservoir properties including nonlinearity, fading memory, state-dependent processing, separability, scalability, and temporal robustness. The neuronal reservoir exhibits nonlinear transformation, achieves 95.83% XOR accuracy at the selected operating point, and retains stimulation-induced state information with characteristic relaxation times of approximately 30–46 ms. Importantly, separability depends on the pre-stimulation network condition, with a near-chaotic regime supporting broader high-separability regions than synchronized activity. Distinct stimulation conditions remain discriminable as the input space increases from 10 to 40 classes, while separability persists despite multi-hour neural drift. Finally, the reservoir is incorporated into a closed perception–action loop for Mario Kart control using a fixed readout. These results establish neuronal cultures as state-dependent living reservoirs whose native dynamics support computation.

## 1 Introduction

Biological neural networks are intrinsically dynamical computing systems whose collective behavior emerges from nonlinear membrane excitability, recurrent synaptic connectivity, excitation–inhibition interactions, synaptic plasticity, and continuously evolving population activity [1–3]. Their responses therefore cannot generally be reduced to static input–output mappings: the same perturbation can propagate differently depending on the instantaneous network condition, while recurrent interactions allow recent inputs to persist transiently in the present population state. Information is distributed across firing activity, spatial recruitment, temporal coordination, and trajectories through a high-dimensional state space, making living neuronal networks natural candidates for dynamical computation.

Reservoir computing provides a theoretical framework for exploiting these properties. Rather than training all recurrent connections, a reservoir maps an input *u*(*t*) through its intrinsic dynamics onto a high-dimensional state **x**(*t*), from which a comparatively simple readout extracts the desired output [4]. A useful reservoir must simultaneously provide nonlinear transformation, input-state separation, and fading memory [4–6]: distinct inputs or input histories should evolve toward distinguishable states [5, 7], while recent perturbations should persist long enough to provide temporal context but ultimately decay [8, 9]. This imposes a fundamental trade-off: recurrent interactions must preserve input-dependent differences sufficiently for separation, yet remain dissipative enough for dependence on remote inputs and initial conditions to fade. The reservoir therefore acts as a nonlinear dynamical kernel that embeds input histories into high-dimensional trajectories accessible to simple readouts [6, 7]. In biological reservoirs, this transformation is inherently state dependent because ongoing neuronal activity forms part of the initial condition from which each input-driven trajectory evolves [3]; computational capability is therefore determined not only by connectivity, but also by the dynamical regime occupied when an input arrives.

Living neuronal cultures provide a direct physical realization of this framework. Early liquid-state-machine experiments established that cultured cortical networks could transform distinct inputs into separable population responses [5], while George *et al*. showed that neuronal cultures can map low-dimensional, nonlinearly separable inputs into higher-dimensional representations accessible to linear readouts [7]. Dissociated cortical networks were also shown to exhibit stimulus-specific short-term and spatiotemporal memory [8, 9]. These studies established state separation, nonlinear state-space transformation, and transient memory as intrinsic computational properties of living neuronal systems.

Subsequent work increasingly formalized and exploited these dynamics. Yada *et al*. demonstrated a living neuronal reservoir coupled to FORCE-based feedback for embodied robot control [10]. Kubota *et al*. applied information-processing-capacity analysis to cultured neuronal networks [11], and Suwa *et al*. quantified first-order memory and second-order nonlinear capacity in dissociated cortical cultures [12]. Sumi *et al*. further demonstrated pattern classification, short-term memory, spoken-digit processing, and generalization using a biological reservoir with a linear readout [6]. Beyond static characterization, Kagan *et al*. embodied human- and rodent-derived cortical cultures in a simulated Pong environment and demonstrated task-related adaptation under closed-loop feedback [13]; subsequent analysis linked task performance to shifts toward near-critical network dynamics [14]. More recently, Robbins et al. demonstrated goal-directed learning in cortical organoids embodied in a closed-loop pole-balancing task, where reinforcement-learning-selected electrical training signals produced task-dependent improvements through neural plasticity [15]. Cai *et al*. extended biological reservoir computing to a three-dimensional human brain organoid in Brainoware, demonstrating speech recognition, nonlinear equation prediction, nonlinear dynamics, and fading memory [16]. Subsequently, Loeffler *et al*. demonstrated handwritten-digit classification using neuronal cultures and showed that computational performance depends strongly on neural architecture, network dynamics, and the choice of decoding method [17]. Other recent studies have reported temporal information retention [18], training-enhanced pattern recognition [19], optogenetic neuronal reservoirs for obstacle avoidance [20], and feedback-driven learning of complex temporal patterns [21]; complementary simulations further indicate that network topology shapes memory, noise robustness, and resilience [22]. Collectively, this literature establishes that living neuronal systems possess nonlinear transformation, transient memory, high-dimensional state structure, and adaptive dynamics relevant to reservoir computation.

The computational framework considered here is summarized in Fig. 1. Application variables are encoded into electrical perturbations through spatial and temporal stimulation dimensions: electrode mapping determines *where* the network is perturbed, while stimulation parameters determine *how* the perturbation is delivered. The culture acts as the physical reservoir, transforming these inputs through native recurrent dynamics into distributed spatiotemporal population responses. Continuous neural-state awareness tracks the instantaneous dynamical condition of the culture, while channel-adaptive spike detection compensates for heterogeneous extracellular recording conditions during construction of the measured reservoir state. A comparatively simple readout then maps the resulting population representation to application-level outputs. This architecture separates input encoding, biological state transformation, state observation, and readout while retaining the evolving neuronal dynamics as the central computational element.

**Fig. 1.**
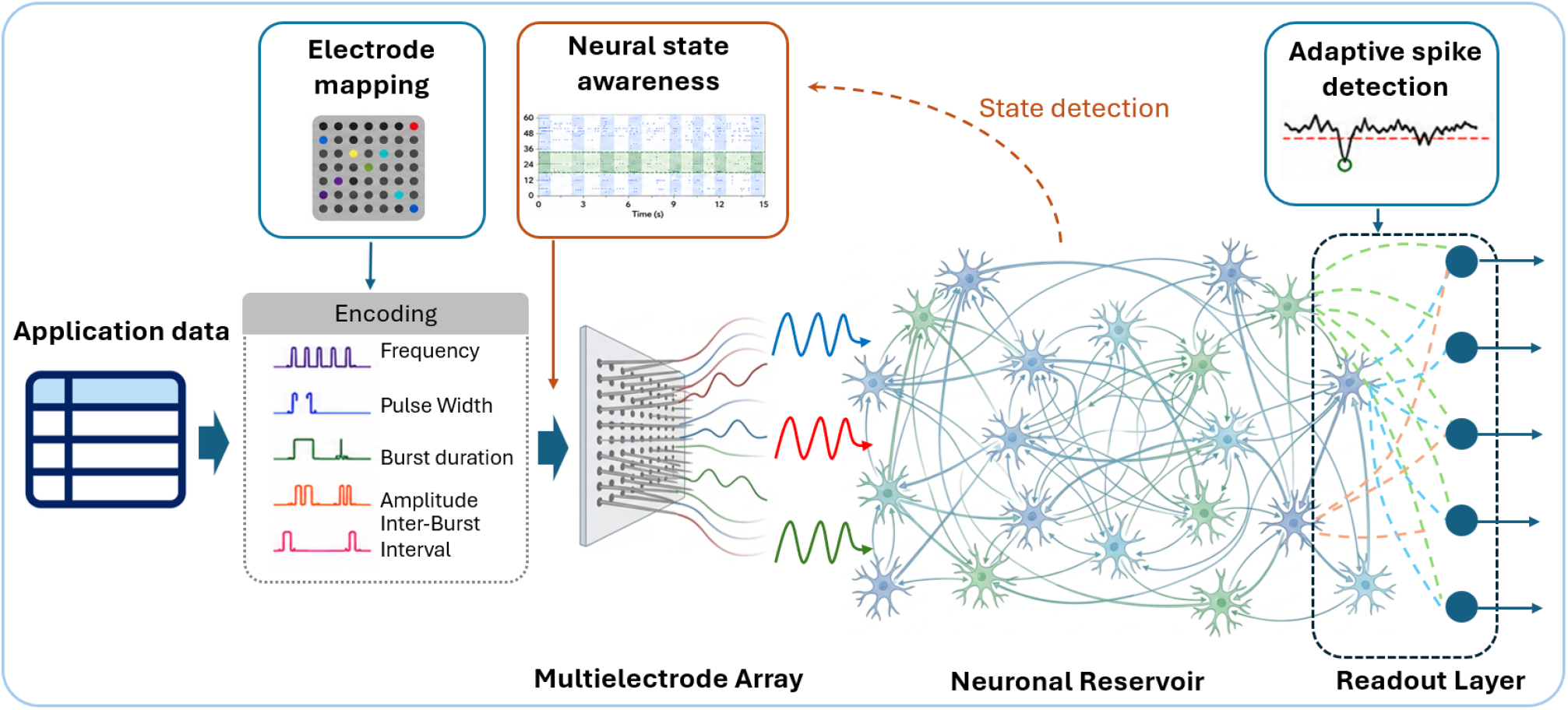
Overview of the neuronal reservoir computing framework. Application data are encoded into electrical stimulation patterns through controllable parameters including stimulation frequency, pulse width, burst duration, amplitude, and inter-burst interval. Candidate stimulation sites are selected through electrode mapping, and the resulting encoded inputs are delivered to the neuronal culture through a multielectrode array (MEA), where the recurrent neuronal network transforms the perturbations into distributed spatiotemporal population responses. Adaptive spike detection provides channel-specific extraction of neuronal events from heterogeneous extracellular recordings, while continuous neural-state awareness characterizes the instantaneous dynamical regime of the reservoir and enables state-informed stimulation. The resulting multichannel neuronal activity forms the reservoir state, which is subsequently projected through a readout layer to generate application-level outputs.

A remaining challenge is that demonstrating a task does not by itself reveal the operating principles that make a living reservoir useful. Unlike fixed artificial reservoirs, neuronal cultures are spatially heterogeneous and intrinsically nonstationary: stimulation and recording sites sample different local populations, spontaneous activity moves the network among distinct dynamical regimes, and excitability, functional connectivity, and population representations evolve over longer timescales. Measurement and analysis can further influence the apparent computational properties of the reservoir, because channel-dependent noise, event-detection sensitivity, and decoding methodology can alter the observed population representation and its measured separability [17]. Previous studies have established individual capabilities including separation, memory, information-processing capacity, classification, adaptation, and embodied control [5–8, 10, 12, 13, 16], but these have largely been examined under different systems and operating conditions. Determining when the *native*, continuously evolving dynamics of a living neuronal network constitute a useful reservoir therefore requires spatial heterogeneity, measurement robustness, nonlinear transformation, memory, instantaneous network state, input-space scaling, and biological drift to be considered together.

Here, we investigate neuronal cultures from this dynamical-systems perspective without deliberately engineering the recurrent biological network using the CL1 MEA systems through the Cortical Cloud [23]. We characterize the unstimulated reservoir by quantifying spatial heterogeneity, spontaneous transitions among population regimes, and longer-timescale reorganization of the state space; introduce a channel-adaptive, median-absolute-deviation-based spike-detection framework to reduce electrode-dependent measurement variability; and directly test nonlinear transformation, XOR processing, and fading memory. To determine whether computation depends on instantaneous network condition, spontaneous activity is classified into random/sparse, synchronized, and an operationally defined near-chaotic regime, and stimulation-evoked separability is compared across states.

Finally, we test whether useful computation persists as both input complexity and biological timescale increase. Stimulation sites are selected through repeated response-based screening accounting for response magnitude, reliability, spatial recruitment, and redundancy. We then evaluate spatio-parametric separability as the number of stimulation conditions increases from 10 to 40 and track the input–state transformation during multi-hour biological drift. The reservoir exhibits nonlinear response transformation, supports XOR separation, and retains stimulation-induced state information over characteristic relaxation times of approximately 30–46 ms. Separability depends strongly on the pre-stimulation network condition, with the operational near-chaotic regime supporting broader high-separability regions than synchronized activity, while useful discrimination persists as the input space expands and the underlying population dynamics drift. We further embed the reservoir in a repeated Mario Kart perception–action loop using a fixed trained readout. Together, these experiments position neuronal cultures not simply as biological classifiers or adaptive agents, but as *state-dependent living reservoirs* whose native dynamics can be systematically characterized and exploited for computation.

## 2 Results

To evaluate how effectively a neuronal culture can function as a computational reservoir, we systematically characterize both its intrinsic network dynamics and its response to controlled perturbations. We examine whether the native neuronal dynamics provide the core properties required for reservoir computing, including rich state-space dynamics, nonlinear information processing, fading memory, input separability, robustness, and scalability. Together, these analyses establish the extent to which computation can be harnessed directly from the culture’s naturally evolving dynamics without modifying the underlying biological network.

### 2.1 Natural dynamics of the neuronal reservoir

Neuronal cultures should not be regarded as spatially uniform or static computing substrates. Even in the absence of externally applied stimulation, the electrophysiological properties of the culture are strongly spatially heterogeneous, such that different regions of the network exhibit distinct local dynamical properties. Fig. 2a shows simultaneous extracellular voltage recordings from two representative MEA channels, Ch. 16 and Ch. 28, selected to highlight distinct channel-specific electrophysiological dynamics over a 0.5-s interval (20.0–20.5 s). Ch. 16 exhibits substantially larger and more frequent voltage excursions, including pronounced negative deflections, whereas Ch. 28 shows comparatively smaller-amplitude fluctuations around its baseline. These contrasting behaviors illustrate that individual recording locations do not constitute equivalent computational nodes; rather, each electrode samples a locally distinct neuronal population with different intrinsic activity characteristics and coupling to the surrounding network.

**Fig. 2.**
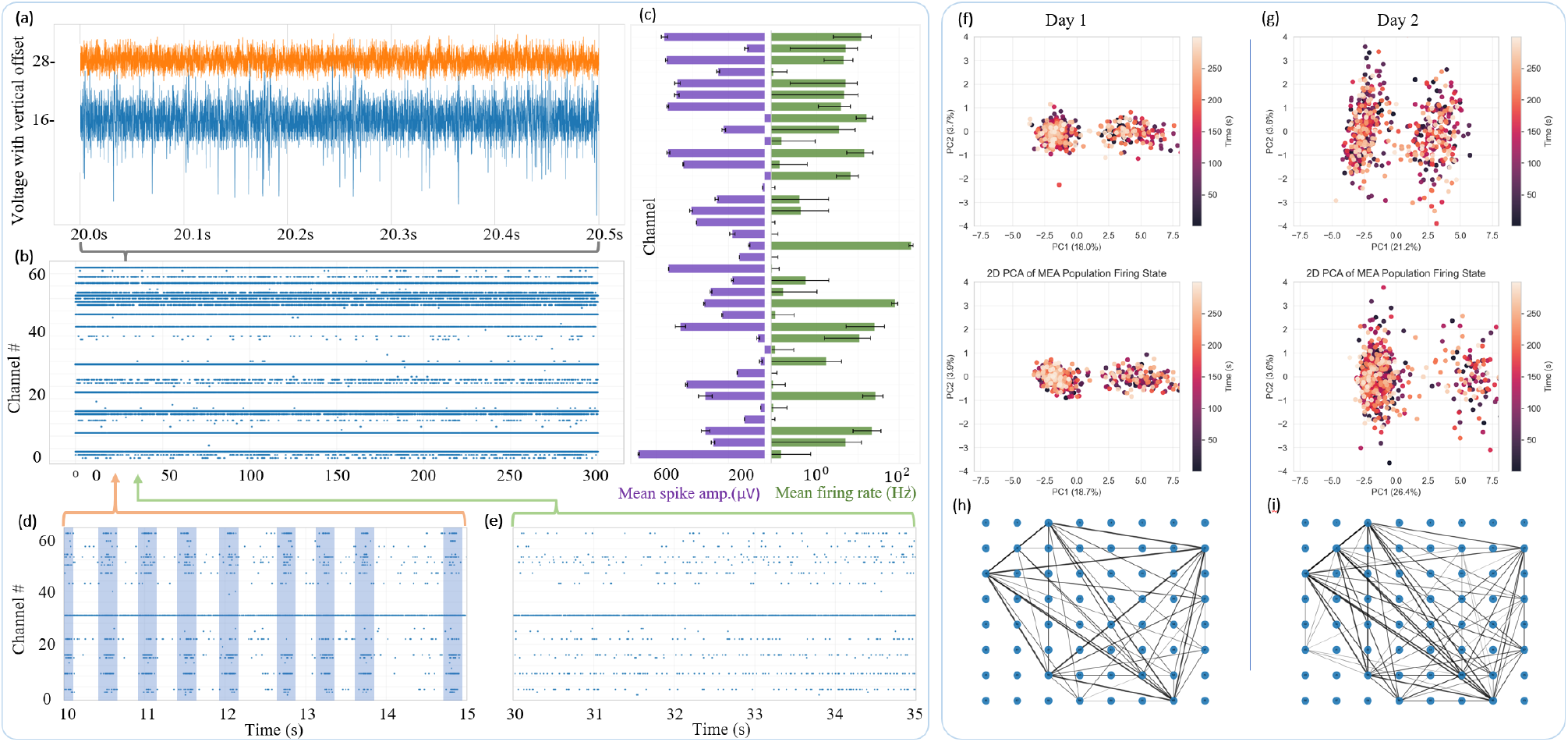
Natural dynamics of the neuronal reservoir under unstimulated conditions. **(a)** Representative extracellular voltage traces recorded from two example microelectrode array (MEA) channels, shown with vertical offsets to illustrate ongoing spontaneous electrophysiological activity at the single-channel level. **(b)** Population-wide spike raster over a 300-s resting recording, demonstrating persistent but spatially and temporally heterogeneous spontaneous activity across the MEA. **(c)** Electrode-wise characterization of spontaneous activity, showing the mean spike amplitude (left, purple; *µ*V) and mean firing rate (right, green; Hz) across recording channels. Error bars indicate the standard deviation, revealing substantial spatial heterogeneity in both extracellular spike magnitude and intrinsic firing activity; the firing-rate axis is displayed on a logarithmic scale to accommodate the broad dynamic range across electrodes. **(d,e)** Representative 5-s epochs extracted from the resting recording, illustrating contrasting endogenous population regimes. The coordinated high-participation regime (synchronized) in **(d)** exhibits recurrent population-wide firing events, highlighted by the shaded intervals, whereas **(e)** shows a comparatively sparse, weakly coordinated regime with reduced and spatially heterogeneous neuronal participation. **(f,g)** Low-dimensional representations of spontaneous population activity on Day 1 and Day 2, respectively, obtained using principal component analysis (PCA). Two representative PCA projections are shown for each day, with points colored according to recording time, revealing structured occupation and temporal evolution of the neuronal population state space. **(h,i)** Functional connectivity networks reconstructed from spontaneous neuronal activity on Day 1 and Day 2, respectively, with nodes corresponding to MEA recording sites and edges representing detected functional interactions between channels.

Moving from the voltage-level behavior in Fig. 2a to the corresponding neuronal firing activity, Fig. 2b shows the spontaneous spike distribution across the full MEA. The raster reveals pronounced channel-to-channel differences in firing rate, with some electrodes exhibiting dense and persistent spiking, others showing intermittent activity, and several remaining comparatively weakly active over the same recording period. Thus, spontaneous activity is distributed nonuniformly across the culture, establishing a spatially heterogeneous reservoir state even before external stimulation is applied.

This heterogeneity is quantified at the electrode level in Fig. 2c, which compares the mean spike amplitude and mean spontaneous firing rate across recording channels, with error bars representing their temporal variability. Large differences are evident in both quantities: some channels generate relatively large extracellular spike amplitudes while others exhibit substantially smaller events, and firing rates span a broad range across the array. Importantly, spike amplitude and firing rate capture complementary aspects of the local electrophysiological state and need not vary proportionally; a recording site may exhibit large-amplitude events without being among the most frequently active channels, or conversely sustain substantial firing with comparatively smaller extracellular amplitudes. This channel-wise diversity therefore reflects heterogeneity not only in neuronal activity level, but also in the strength and statistical structure of the extracellular activity sampled throughout the network. From a reservoir-computing perspective, such spatial diversity provides a naturally heterogeneous collection of dynamical units from which distributed population responses can emerge.

Beyond this spatial heterogeneity, the neuronal reservoir exhibits rapidly evolving temporal dynamics characterized by spontaneous transitions among distinct network states. In Fig. 2d, the culture enters a strongly synchronized regime, where recurrent population-wide events appear as vertical structures in the raster, reflecting highly coordinated firing across many channels. In contrast, Fig. 2e shows a weakly coordinated regime in which synchronous events are largely absent and firing is more irregularly distributed across space and time. The rapid transition between these regimes demonstrates that the reservoir does not operate around a single stationary condition but continuously explores different dynamical configurations. Between these extremes, the network can occupy an intermediate *near-chaotic* regime that preserves greater response diversity while retaining sufficient recurrent interaction for perturbations to propagate through the reservoir. The extent to which this state-dependent dynamical diversity influences the computational response of the reservoir is investigated explicitly in Section 2.2.3.

The PCA projections in Fig. 2f,g provide a complementary population-level view of the spontaneous reservoir dynamics across multiple timescales. The coexistence of compact and diffuse regions within the projected state space reflects differences in the range of population configurations explored by the culture and, when interpreted alongside the raster patterns, is consistent with the contrasting coordinated and weakly coordinated activity regimes described above. Beyond these rapid state transitions, comparison across days reveals a pronounced reorganization of the spontaneous reservoir state space, providing evidence of neural drift at the population level. Each point represents the instantaneous MEA population activity projected onto the dominant principal components and thus corresponds to a low-dimensional representation of the reservoir state, **x**(*t*). On Day 1 (Fig. 2f), the population activity occupies comparatively compact and confined regions of the projected state space, whereas by Day 2 (Fig. 2g), these regions become substantially broader and redistributed across the principal-component directions. Importantly, this temporal reorganization is also evident in the functional network structure shown in Fig. 2h,i. Here, nodes represent MEA recording locations and edges represent functional relationships between their spontaneous activities. The spatial pattern and distribution of these connections change markedly between Day 1 and Day 2, indicating that the underlying functional coupling among neuronal populations is itself reorganized over time. Thus, the observed drift is not restricted to a change in the low-dimensional projection of population activity, but is accompanied by a corresponding alteration in the functional interaction structure of the neuronal network.

Together, the changes in population-state geometry and functional connectivity are consistent with *neural drift*, whereby the reservoir’s broader activity structure and functional organization evolve over time beyond its rapid switching among transient network states. Such evolution can modify the population trajectories and interaction pathways through which external perturbations are transformed, making the effective reservoir mapping potentially time dependent. The neuronal culture should therefore not be treated as a static, time-invariant computing substrate; its computational characterization must account for both short-timescale state transitions and longer-timescale reorganization of the underlying neuronal dynamics.

### 2.2 Computational dynamics of the neuronal reservoir

Having established that the neuronal culture exhibits intrinsically heterogeneous, nonstationary, and recurrent spontaneous dynamics, we next ask whether these native dynamics can be harnessed as a computational reservoir by testing their capacity for nonlinear transformation, fading memory, state-dependent processing, and robust separation of externally imposed inputs.

#### 2.2.1 Nonlinear Perturbation Dynamics and Fading Memory

A neuronal culture can function as a computational reservoir only if it can nonlinearly transform incoming perturbations while retaining their influence for a finite duration. Nonlinearity enables the network to map inputs into richer population-state representations that support separation of complex input relationships, whereas fading memory allows the current reservoir state to preserve recent temporal context without becoming permanently dominated by past inputs. The computationally useful regime therefore requires a balance between rapid relaxation and persistent state retention. Here, we evaluate these properties by quantifying deviations from linear response superposition, testing nonlinear XOR computation, and measuring the temporal decay of stimulation-induced reservoir states.

To evaluate nonlinear integration, we first performed a fundamental superposition test, as shown in Fig. 3(a), by comparing the measured response to combined stimulation with the linear prediction obtained by summing the responses to the corresponding individual perturbations. For a linear system, these measured and predicted responses should satisfy the identity relation, *R*(*A* + *B*) = *R*(*A*) + *R*(*B*). Instead, systematic deviations from the identity line were observed across recording channels, demonstrating that the neuronal reservoir does not simply superimpose independently evoked responses, but nonlinearly transforms combined inputs. The channel-wise nonlinearity index, defined as the normalized deviation of the observed combined response from the linear superposition prediction (see Methods), revealed substantial heterogeneity across input electrodes as shown in Fig. 3(b), indicating that different regions of the reservoir exhibit different degrees of nonlinear transformation. Together, these results demonstrate distributed and spatially heterogeneous nonlinear processing within the neuronal reservoir, providing an important computational property for expanding input representations into a richer reservoir state space.

**Fig. 3.**
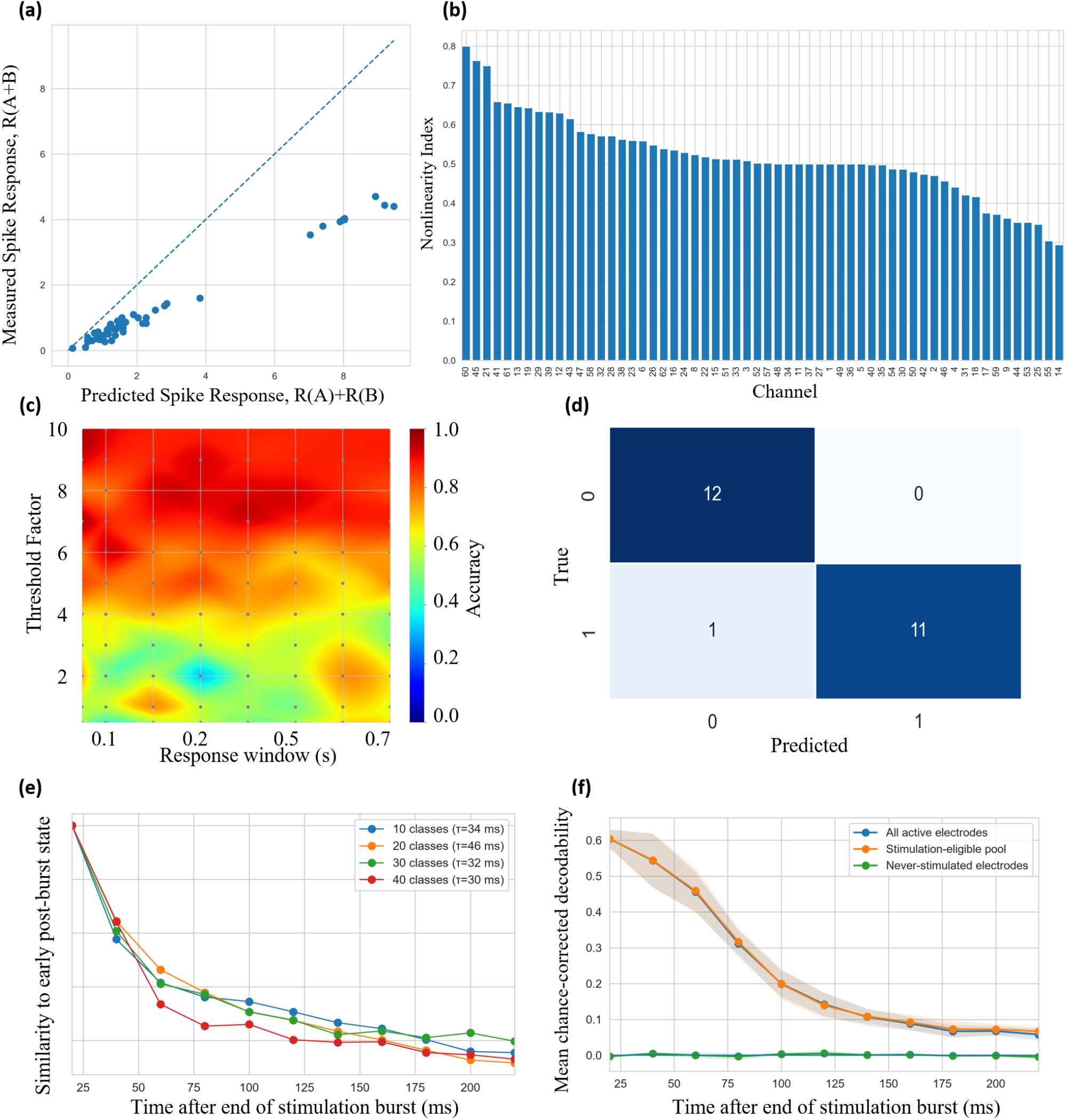
Nonlinear response transformation and fading-memory dynamics of the neuronal reservoir. **(a)** Channel-wise test of linear superposition, comparing the response observed under combined stimulation *A* + *B* with the response predicted from the sum of the independently evoked responses to *A* and *B*. The dashed identity line represents the expectation for a purely linear system, such that systematic deviations quantify nonlinear integration. **(b)** Channel-wise nonlinearity index, revealing heterogeneous nonlinear transformation across the electrode population. **(c)** XOR readout accuracy as a function of the post-stimulation response window and spike-detection threshold factor. The broad region of high accuracy demonstrates that nonlinear computational performance is preserved across a range of response-extraction parameters rather than being restricted to a narrowly tuned operating point. **(d)** Confusion matrix for the XOR readout at the selected operating point (50 ms response window, threshold factor 7.0), yielding an accuracy of 0.9583 (23*/*24 correctly decoded trials). Successful XOR decoding provides a functional demonstration that the neuronal reservoir transforms the input into a representation supporting nonlinear separation. **(e)** Fading of the stimulation-induced population representation, quantified as the similarity between the evolving post-burst reservoir state and the early post-burst state for experiments containing 10, 20, 30, and 40 distinct input conditions. The exponential-like decay yields characteristic fading times of approximately *τ* = 34, 46, 32, and 30 ms, respectively, demonstrating transient retention followed by progressive loss of stimulus-related state information. **(f)** Spatial control analysis of the fading-memory trace, showing comparable time-dependent decodability when using all active electrodes or only the stimulation-eligible electrode pool, whereas never-stimulated electrodes remain near chance level.

To test whether the observed nonlinear transformations were computationally useful rather than random deviations from linearity, we evaluated the reservoir using the XOR task, a canonical nonlinear benchmark that cannot be solved by a linear mapping in the original input space. Reservoir performance was first examined across combinations of post-stimulation response windows and spike-detection thresholds [Fig. 3(c)]. High XOR performance was maintained across a broad region of the parameter space, indicating that the nonlinear computation was not restricted to a narrowly tuned analysis condition. At the selected operating point, corresponding to a 50 ms response window and threshold factor of 7, the reservoir achieved an XOR accuracy of 95.83% as shown in Fig. 3(d). Because the readout operates on the neuronal population response, successful XOR discrimination indicates that the recurrent network transformed the originally linearly inseparable input relationships into a higher-dimensional reservoir representation in which the output classes became separable. Thus, the physical response nonlinearity observed in Fig. 3(a,b) is accompanied by a functional nonlinear transformation capable of supporting computation.

Next, we evaluated fading memory by quantifying the persistence and relaxation of stimulation-induced reservoir states across post-stimulation delays and electrode populations as shown in Fig. 3(e,f). The normalized perturbation memory, *M* (*t*), progressively decayed following stimulation, yielding characteristic relaxation timescales of approximately *τ* = 34, 46, 32, and 30 ms for the 10-, 20-, 30-, and 40-input sets, respectively. The similarity of the evolving population state to the early post-stimulation state likewise decreased with time, indicating progressive loss of the initial stimulus-evoked representation. The comparable relaxation timescales across input-set sizes suggest that the fading dynamics are primarily governed by the intrinsic relaxation properties of the neuronal reservoir rather than by the number of encoded inputs. Spatially, the all-active and stimulation-eligible electrode populations exhibited similar time-dependent decodability, whereas never-stimulated electrodes remained near chance level [Fig. 3(f)], localizing the retained input information predominantly to the stimulation-responsive network. Together, these results demonstrate that external perturbations generate distributed reservoir states whose magnitude and representational structure persist transiently before progressively relaxing, providing the finite temporal dependence required for fading-memory computation.

#### 2.2.3 Adaptive spike detection performance

Reliable characterization of the neuronal reservoir requires transforming continuous extracellular voltages into discrete neuronal events. As established in Section 2.1, the culture exhibits pronounced spatial heterogeneity, with substantial channel-to-channel variation in baseline amplitude and activity. Such variability, together with temporal non-stationarity, makes a fixed global threshold susceptible to channel-dependent detection bias. We, therefore, employed a channel-adaptive spike-detection framework that scales the detection criterion to the local statistical background of each electrode.

Figure 4a presents the proposed channel-adaptive spike-detection algorithm used to extract neuronal events from the multichannel extracellular recordings. For each MEA channel, the local noise level is estimated independently using a robust median absolute deviation (MAD)-based statistic, from which a channel-specific negative detection threshold is derived. Threshold crossings are further constrained by a refractory period to prevent multiple detections of the same extracellular event. The resulting spike times are subsequently used to construct the channel-wise activity features that define the reservoir state. A complete description of the detection procedure and parameterization is provided in the Methods section.

**Fig. 4.**
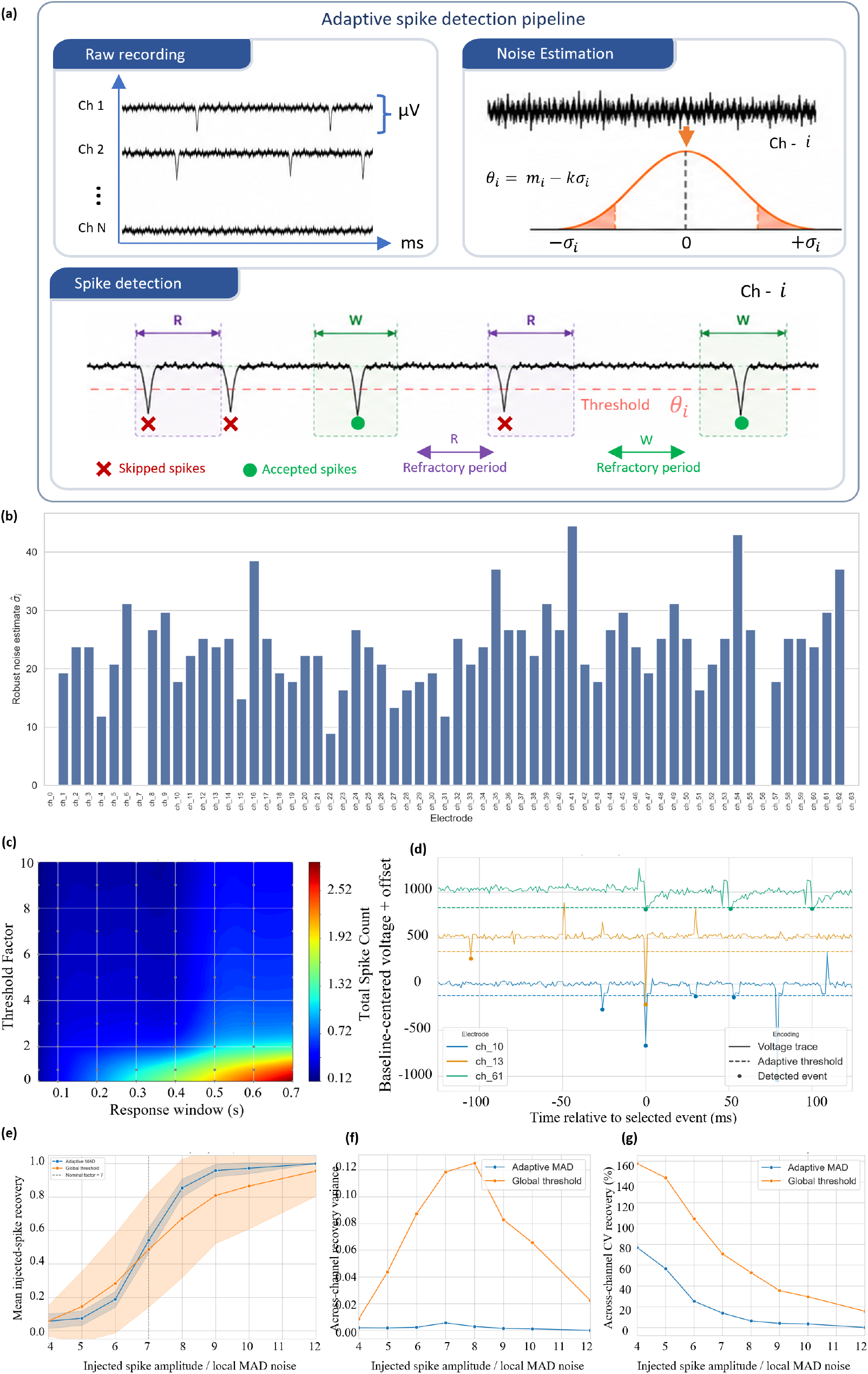
Adaptive spike detection across heterogeneous MEA recording channels. (a) Schematic of the adaptive spike-detection pipeline, comprising raw multichannel acquisition, robust channel-wise noise estimation, adaptive threshold crossing, refractory-period enforcement, and response-window spike counting. (b) Channel-specific robust background-noise estimates across the MEA, demonstrating substantial spatial variability in the local electrophysiological recording conditions. (c) Sensitivity of the total detected spike count to the threshold factor and response-window duration, illustrating the dependence of event yield on detector stringency and temporal integration. (d) Representative baseline-centered voltage traces from three electrodes with independently estimated adaptive thresholds and detected spike events, demonstrating channel-specific normalization of the detection criterion. (e) Mean recovery of synthetically injected spikes as a function of spike amplitude normalized by the local MAD noise level, comparing the adaptive MAD detector with a global-threshold approach; the dashed line indicates the nominal threshold factor of 7. (f) Across-channel variance in injected-spike recovery, showing substantially lower electrode-dependent variability for the adaptive detector. (g) Corresponding coefficient of variation of recovery across channels, further demonstrating improved consistency of adaptive detection across heterogeneous recording sites. Together, these results show that channel-wise MAD normalization provides a more uniform and robust event-detection interface for constructing neuronal reservoir states from spatially heterogeneous extracellular recordings.

Fig. 4b shows that the robust background-noise scale varies markedly across the MEA, spanning approximately 8–45 *µ*V across active electrodes. This more than fivefold variation indicates that an identical absolute voltage excursion would not represent an equivalent event across recording sites, making a single global voltage threshold intrinsically biased toward particular electrodes.

The parameter dependence of event extraction is evident in Fig. 4c. At high threshold factors, detected event yield remains low across response windows, whereas progressively more events are admitted as the threshold is reduced and the temporal integration window is increased. The strongest increase occurs at low threshold factors and longer response windows, demonstrating that both detection stringency and temporal integration directly shape the observable reservoir state. Figure 4d provides representative traces from three electrodes with markedly different baseline fluctuation scales; despite these differences, the adaptive detector assigns channel-specific thresholds that preserve a common statistical detection criterion across recording sites.

A reliable multichannel reservoir state requires equivalent neuronal events to be detected consistently across electrodes despite differences in their background signal levels. Therefore, we tested whether the adaptive detector reduces electrode-dependent detection bias by superimposing synthetic extracellular spike waveforms of known timing and amplitude onto measured backgrounds from electrodes spanning the observed noise distribution. This provided a controlled ground truth for comparing adaptive and global thresholding. Figure 4e shows that the injected-spike detection rate increased with normalized spike amplitude for both methods, but adaptive thresholding reached high detection rates more rapidly. Near the selected threshold factor of 7, the adaptive detector identified approximately 60–70% of injected events, compared with approximately 40–50% for global thresholding. The substantially wider dispersion under global thresholding reflects its use of a common absolute detection boundary across channels with different local noise scales, causing equivalent normalized events to have different effective detection sensitivities across electrodes. In contrast, the adaptive detector scales the threshold to each channel’s local MAD noise and therefore maintains a more consistent detection criterion. The effect of channel-wise normalization is further quantified in Fig. 4f,g by examining the dispersion of spike-recovery performance across electrodes. Figure 4f shows the across-channel variance in recovery as a function of injected-spike amplitude relative to the local MAD noise level. The adaptive method maintains a consistently low variance across nearly the entire amplitude range, indicating that electrodes with different intrinsic noise levels and signal amplitudes achieve comparable recovery performance after local normalization. In contrast, the global-threshold method exhibits a pronounced increase in variance at intermediate signal-to-noise ratios, with maximal dispersion near the detection-transition regime. This behavior indicates that a single global threshold causes different electrodes to enter the detectable regime at substantially different effective signal strengths, producing strong channel-dependent detection bias.

Figure 4g provides a complementary normalized measure through the across-channel coefficient of variation (CV) of spike recovery. The adaptive method produces a markedly lower CV across the tested amplitude range, demonstrating that the reduction in channel-to-channel dispersion persists even after accounting for differences in the mean recovery level. The contrast is particularly strong at low-to-intermediate relative spike amplitudes, where detection is most sensitive to local noise statistics. As the injected spikes become increasingly distinguishable from the background, the CV decreases for both methods because recovery approaches saturation across electrodes; nevertheless, adaptive MAD normalization remains substantially more consistent. Together, panels (f) and (g) demonstrate that channel-adaptive thresholding not only improves mean spike recovery but, more importantly, reduces electrode-dependent variability in event detection, thereby producing a more uniform basis for constructing the population reservoir state. These results demonstrate that adaptive thresholding reduces channel-dependent detection bias, providing a more uniform multichannel event representation for constructing the neuronal reservoir state.

Together, these results show that the principal advantage of channel-adaptive detection is not simply higher spike recovery, but substantially improved consistency across heterogeneous electrodes. This is critical for reservoir analysis because each electrode contributes a dimension to the population state vector; unequal detection sensitivity would therefore distort the geometry of the measured reservoir state before any computational property is assessed. Adaptive MAD normalization provides a more statistically uniform observation layer across the MEA, reducing the likelihood that subsequent differences in nonlinear transformation, fading memory, state dependence, or input separability arise from channel-dependent measurement bias.

#### 2.2.3 State-dependent input processing in the neuronal reservoir

The computational response of a neuronal reservoir is not determined solely by the external input, but also by the dynamical state of the network at the moment the perturbation is applied. In a recurrent biological network, spontaneous activity continuously reshapes membrane excitability, synaptic efficacy, population synchrony, and the distribution of active neuronal subpopulations, such that the effective input–state transformation is intrinsically time dependent. Formally, the evoked reservoir state can be expressed as

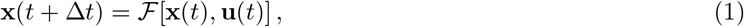

where the same input **u**(*t*) may generate different trajectories depending on the pre-stimulation state **x**(*t*). Consequently, spontaneous transitions among sparse, near-chaotic, and synchronized regimes should be interpreted as transitions among distinct computational operating conditions rather than as background variability. These regimes differ in the balance between collective coherence and dynamical freedom: excessive synchronization can constrain the accessible state space and compress response diversity, whereas a more weakly coordinated, near-chaotic regime can preserve sensitivity to perturbations while allowing inputs to evolve into distinct population trajectories. We therefore characterize the spontaneous state structure of the neuronal culture and directly test how stimulation delivered in different dynamical regimes alters the separability of the resulting reservoir responses, establishing whether computational capability is intrinsically state dependent.

The synchronization index in Fig. 5a varied substantially over the 300-s recording, while the active-electrode fraction and population spike count in Fig. 5b,c showed repeated changes in spatial recruitment and overall firing intensity. Taken together, these observables were used to operationally classify the instantaneous reservoir condition into random/sparse, near-chaotic, and synchronized regimes, shown by the color-coded state trajectory beneath Fig. 5c. The frequent transitions among these states demonstrate an intrinsically nonstationary and metastable network organization in which the dynamical context of the reservoir evolves on the timescale of seconds.

**Fig. 5.**
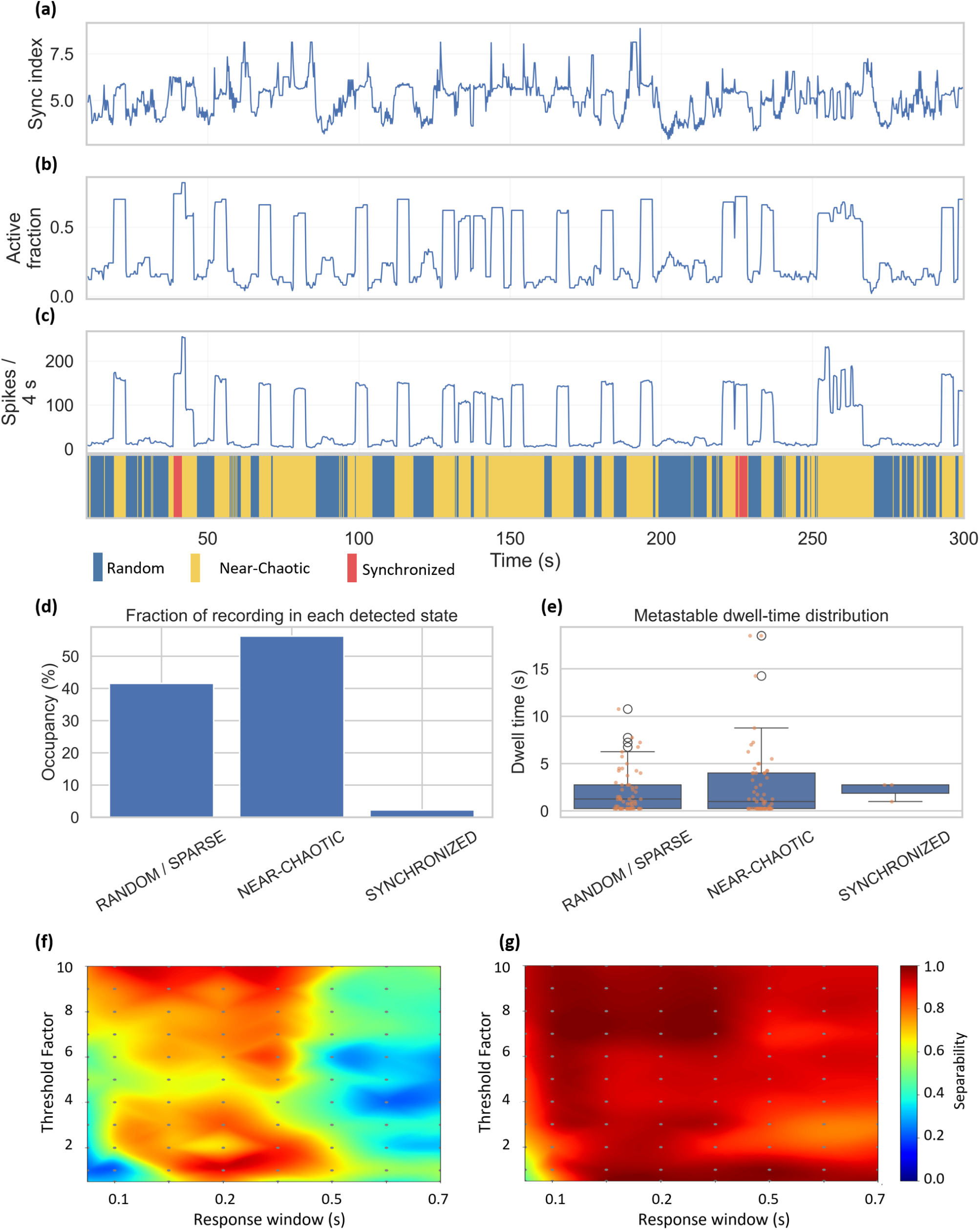
State-aware characterization of neuronal dynamics and state-dependent input separability. (a–c) Temporal evolution of the network synchronization index, fraction of active recording channels, and population spike count, respectively, computed over successive 4-s analysis windows across the 300-s recording. The color-coded state trajectory below (c) shows the corresponding classification of the instantaneous reservoir dynamics into random/sparse, near-chaotic, and synchronized regimes. (d) Fraction of the recording occupied by each detected state, showing predominant occupancy of the near-chaotic regime and only transient occurrence of synchronized activity. (e) Distribution of metastable dwell times for each detected state, quantifying the temporal persistence of the corresponding dynamical regimes; individual dwell intervals are overlaid on the distributions. (f) Spatio-parametric input-separability landscape for stimulation delivered during the synchronized state as a function of spike-detection threshold factor and post-stimulation response window, revealing strong dependence of separability on the response-extraction parameters. (g) Corresponding separability landscape for stimulation delivered during the near-chaotic state, showing consistently high separability across a broad range of threshold factors and response windows. Color in (f,g) denotes the normalized separability score, with values approaching unity indicating greater differentiation among stimulation-evoked reservoir responses.

The temporal organization of these regimes is summarized in Fig. 5d,e. The culture occupied the nearchaotic regime most frequently (51.3%), followed by the random/sparse regime (43.9%), whereas synchronized activity accounted for only 4.8% of the recording. The corresponding dwell-time distributions show that these states persisted for finite intervals before transitioning, supporting their interpretation as metastable network configurations rather than isolated fluctuations. This state occupancy is computationally relevant because each regime defines a different balance of synchrony, neuronal recruitment, and recurrent interaction at the time an external perturbation is applied.

The effect of the pre-stimulation network state is evident in the separability landscapes of Fig. 5f,g. Stimulation delivered during synchronized activity produced a strongly parameter-dependent response landscape, with separability varying substantially across threshold factors and response windows. In contrast, stimulation during the near-chaotic regime yielded consistently high separability over a much broader region of the analysis space. This indicates that near-chaotic activity provides a more favorable operating condition for reservoir computation, preserving sufficient dynamical diversity for distinct inputs to evolve into distinguishable population responses while avoiding the strong state-space compression associated with highly synchronized activity. Together, these results demonstrate that the computational response of the neuronal reservoir is intrinsically state dependent and identify the near-chaotic regime as a robust operating condition for input separation.

#### 2.2.4 Robustness, Scalability, and Reproducibility of Spatio-Parametric Input Separability

In this section, we focus primarily on reservoir performance in terms of state separability, a central requirement for reservoir computing because distinct inputs must be transformed into sufficiently differentiated internal representations to support reliable downstream readout. Before evaluating separability, however, the input electrodes themselves must be appropriately selected. As demonstrated by the intrinsic spatial heterogeneity of the culture in Section 1, stimulation at different electrodes does not perturb equivalent neuronal populations and can therefore differ substantially in the magnitude, reliability, spatial recruitment, and population pattern of the resulting response. We therefore performed systematic *stimulation-electrode screening*, in which each candidate electrode was repeatedly stimulated under an identical perturbation and ranked using a composite measure of evoked-response magnitude, response reliability, and spatial recruitment, while additionally excluding highly correlated response maps to preserve diversity among the selected stimulation sites (Methods).

Because the neuronal substrate is intrinsically variable, electrode selection based on a single perturbation could reflect trial-specific fluctuations rather than a reproducible property of the stimulation site. We therefore first examined how the screening estimates stabilized with repeated sampling. Figure 6a shows that the coefficient of variation of the screening score decreased progressively as additional repetitions were incorporated, whereas the corresponding evoked-response variability approached a comparatively stable level. This indicates that repeated sampling progressively stabilizes the composite estimate used to rank stimulation sites despite persistent biological variability in the underlying evoked responses. Consistent with this interpretation, Fig. 6b shows that the relative change in screening score was initially large but rapidly decreased with successive repetitions, approaching only small incremental changes at higher repeat numbers. Notably, this convergence is largely achieved by approximately *∼*10 screening repetitions, beyond which additional repeats provide only marginal improvement in score stability, suggesting that ~10 repeats are sufficient for reliable stimulation-site ranking under these conditions. Thus, additional trials progressively contributed less information to the estimated electrode ranking, indicating convergence of the screening procedure.

**Fig. 6.**
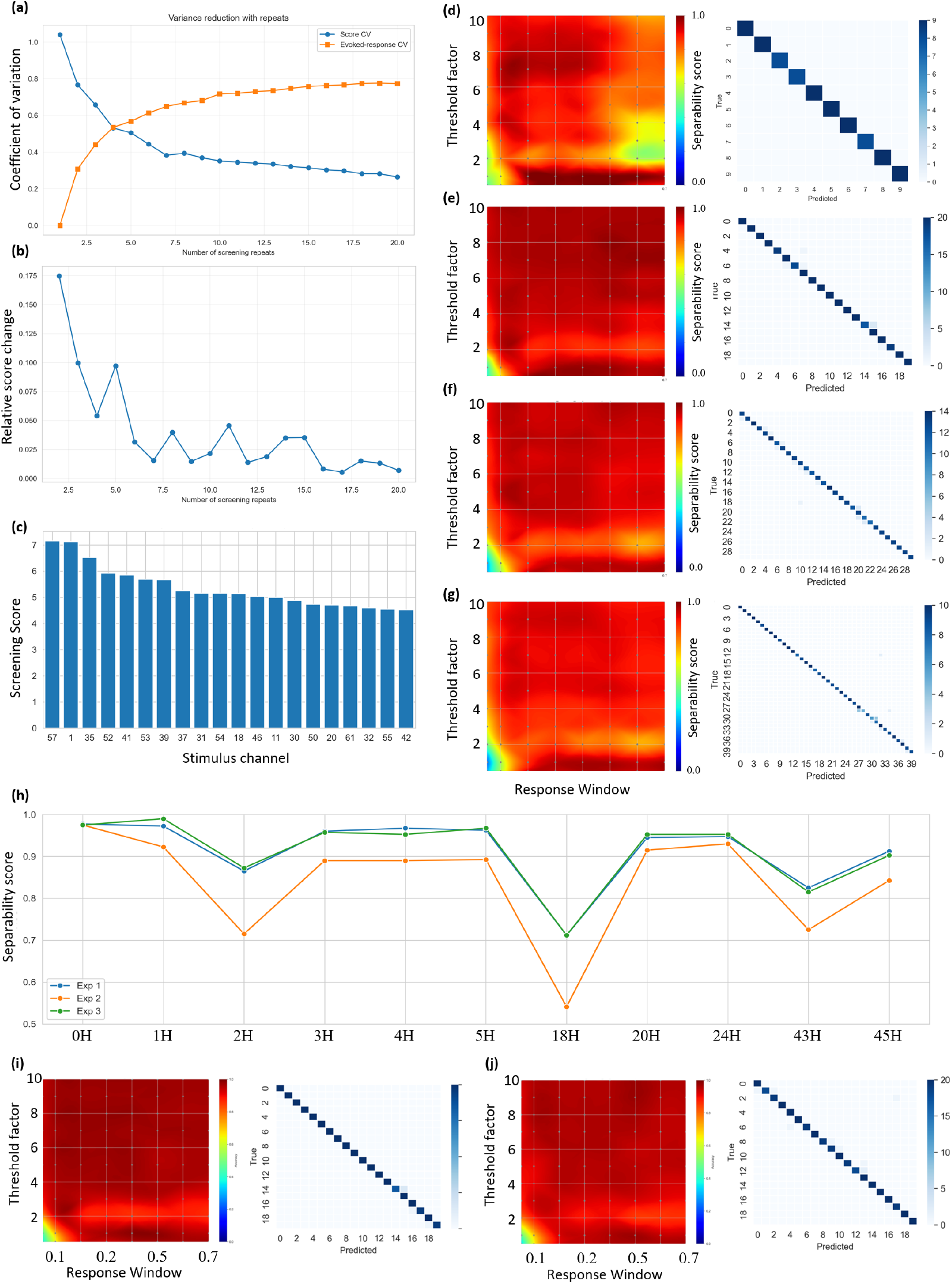
Robustness, scalability, and temporal stability of spatio-parametric input separability in the neuronal reservoir. (a) Dependence of stimulation-channel screening variability on the number of repeated trials, quantified using the coefficient of variation (CV) of the screening score and stimulation-evoked response magnitude. The progressive reduction in screening-score variability indicates increasing reliability of the estimated channel response with repeated sampling. (b) Marginal information gain contributed by each additional screening repetition, quantified from the relative change in screening score. The rapid decay toward small incremental changes indicates convergence of the channel characterization and provides an empirical basis for selecting a sufficient number of screening repeats. (c) Resulting stimulation-electrode ranking obtained after the screening procedure, showing substantial spatial heterogeneity in channel-specific screening scores and identifying the electrodes that most reliably generate informative reservoir responses. (d–g) Separability landscapes and corresponding confusion matrices for progressively increasing spatio-parametric input sets containing 10, 20, 30, and 40 distinct stimulation conditions, respectively. Separability is evaluated across combinations of spike-detection threshold factor and post-stimulation response window. The broad regions of high separability and strongly diagonal confusion matrices demonstrate that distinct stimulation conditions remain discriminable as the cardinality of the input space increases. (h) Temporal evolution of 10-class readout accuracy measured at representative stimulation channels across three experiments over approximately 45 h, revealing the effect of biological neural drift on the separability of stimulation-evoked reservoir representations. Although the magnitude of drift is channel- and experiment-dependent, substantial input separability is retained over the recording interval. (i,j) Representative 20-class separability landscapes and corresponding confusion matrices obtained approximately 20 h apart, illustrating the temporal evolution of the reservoir input–response mapping under neural drift. Despite changes in the underlying biological state, both measurements retain broad high-separability regions and predominantly diagonal confusion structure. Color in the separability landscapes denotes the normalized separability score, with values approaching unity indicating stronger differentiation among stimulation-evoked reservoir states. Collectively, these results demonstrate that reliable stimulation electrodes can be identified through convergence-based repeated screening, that the neuronal reservoir preserves separability as the number of spatio-parametric input conditions increases, and that useful input discrimination persists despite biological drift over extended experimental timescales.

The converged screening scores revealed substantial differences in stimulation efficacy across the candidate electrodes, as shown in Fig. 6c. Rather than forming an approximately uniform distribution, the ranked scores span a broad range, with a subset of electrodes producing substantially stronger and more reliable distributed responses than others. Together, Fig. 6a–c establishes that stimulation location constitutes an important input dimension of the neuronal reservoir and that informative stimulation sites can be identified reproducibly through repeated, response-based screening. The resulting stimulation-electrode set was subsequently used to evaluate whether distinct spatio-parametric perturbations remain separable as the complexity of the input space increases.

The scalability of the neuronal reservoir was evaluated by progressively increasing the number of spatio-parametric input conditions from 10 to 40 classes and examining both the separability landscape and the corresponding class-confusion structure, as shown in Fig. 6(d–g). Across all input-set sizes, high separability was maintained over a broad region of the response-extraction parameter space, particularly at moderate-to-high spike-detection thresholds and response windows sufficiently long to capture the distributed post-stimulation dynamics. This persistence is important because it indicates that the underlying reservoir representation does not collapse as the number of distinct input conditions increases; instead, the network continues to map different stimulation configurations onto distinguishable population states despite the increasing density of representations in state space. The corresponding confusion matrices remained strongly concentrated along the diagonal for 10-, 20-, 30-, and 40-class evaluations, demonstrating that the evoked responses preserved class-specific structure with comparatively limited cross-class overlap. From a dynamical-systems perspective, these results indicate that the neuronal reservoir supports a scalable input–state transformation in which additional spatio-parametric perturbations occupy distinct regions of the accessible response manifold rather than converging onto a small set of redundant trajectories. The broad high-separability regions further show that this scaling behavior is not dependent on a narrowly optimized response window or spike-detection threshold, supporting the interpretation that the observed discrimination reflects intrinsic structure in the stimulation-evoked population dynamics rather than an artifact of a particular analysis setting.

The temporal stability of the neuronal reservoir was evaluated by tracking input separability over extended recording periods, thereby probing the effect of biological neural drift on the stimulation–response mapping. Figure 6(h) shows the 10-class separability accuracy measured across representative stimulation channels in three independent experiments over approximately 45 h. Although the absolute performance exhibited channel- and experiment-specific fluctuations, reflecting the gradual reconfiguration of excitability, connectivity, and spontaneous network state expected in a living neuronal substrate, the overall separability remained high for most channels throughout the recording period. This indicates that neural drift perturbs the detailed geometry of the reservoir response space without necessarily destroying the computational distinction among input conditions. The corresponding separability landscapes acquired approximately 20 h apart, shown in Fig. 6(i,j), provide a direct visualization of this temporal evolution. Despite measurable changes in the response landscape, both measurements retained broad regions of high separability and strongly diagonal confusion matrices, demonstrating that the population representation remained functionally discriminative across substantial biological drift. Together, these results show that the neuronal reservoir is not temporally invariant, but its input–state mapping possesses sufficient dynamical redundancy and distributed encoding to preserve useful separability over multi-hour timescales.

### 2.3 Game-Play Steering Control Using a Neuronal Reservoir

As a real-time application demonstration, the neuronal reservoir was evaluated in a closed perception–action loop for *Mario Kart Wii* steering control, as illustrated in Fig. 7. Figure 7(a) shows the game environment during closed-loop operation, including the kart, track, and surrounding track features. On the host PC, the game state was read directly from memory to obtain the kart position and orientation. These data were projected onto a previously defined track centerline, from which the nearest centerline position and a forward lookahead point were identified, as illustrated in Fig. 7(b).

**Fig. 7.**
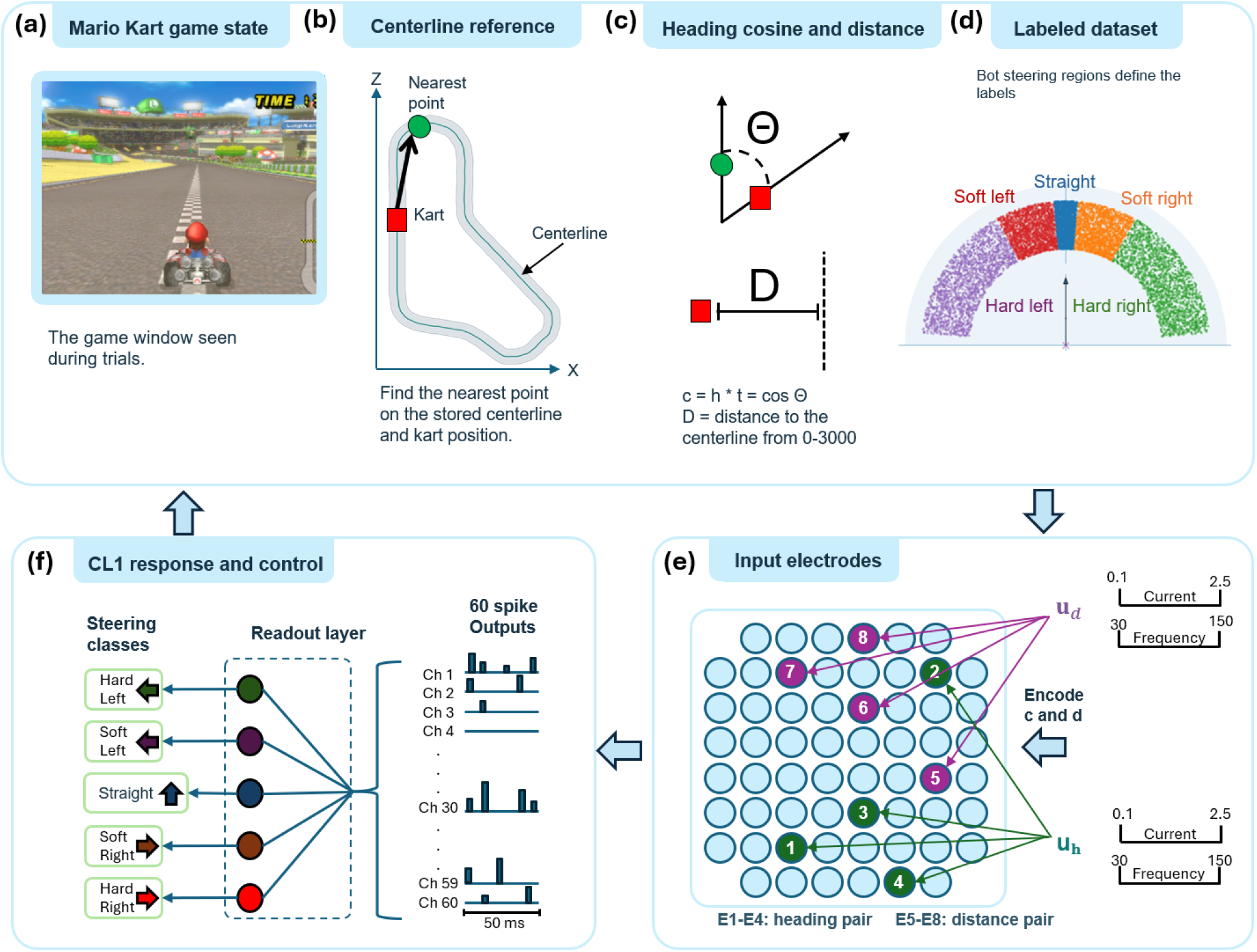
Closed-loop neuronal-reservoir steering pipeline. (a) The game state is read from memory. (b) Kart position is projected onto a stored track centerline, and a lookahead point defines the desired heading. (c) The client computes the signed heading error, *θ*; centerline distance is retained for tracking analysis only and is not encoded as a neuronal input. (d) The same angle is discretised into five reference steering classes for performance evaluation; the reference class is never substituted for a neuronal prediction. (e) The scalar *u* = clip(*θ/*45°, *−*1, 1) jointly determines the stimulation frequency, *f* = 105 + 75*u* Hz, and current, *I* = 1.3 + 1.2*u µ*A on four electrodes while the distance from the centerline is equally mapped in a similar way to stimulate 4 electrodes. A three-pulse, charge-balanced biphasic stimulus is then delivered through all eight selected electrodes. (f) The resulting multichannel neuronal response is converted into the specified 60-feature vector and evaluated by the fixed pretrained ridge readout, which returns one of five steering actions to the game client.

The geometric relationship between the kart and the desired trajectory was then calculated, as shown in Fig. 7(c). A signed heading error was determined from the kart heading vector and the vector directed toward the selected centerline lookahead point. In addition, the distance between the kart and the track centerline was calculated to quantify the kart’s displacement from the desired trajectory. Together, the signed heading error and centerline distance provided the continuous state variables used to encode the current game condition.

For evaluation purposes, the current game state was also assigned to one of five reference steering classes, as shown in Fig. 7(d). These reference classes represented the steering action predicted by the control policy and were used as ground-truth labels for evaluating neuronal-reservoir performance. During closed-loop operation, however, the reference class was never substituted for the neuronal prediction. This labeled data was also used as a dataset for training, with 800 of each class in the dataset, 4000 total data samples.

The signed heading error and centerline distance were subsequently mapped to the electrical stimulation parameters delivered to the neuronal culture, as illustrated in Fig. 7(e). These continuous state variables were encoded across a stimulation-frequency range of 30–180 Hz and a current range of 0.1–2.5 *µ*A, respectively. Stimulation was delivered through eight selected electrodes, with four electrodes assigned to each encoded variable. An identical charge-balanced, three-pulse biphasic stimulation pattern was applied within each electrode group according to the corresponding encoded value. The stimulation electrodes were selected using the candidate-search procedure described previously.

Finally, as shown in Fig. 7(f), the resulting multichannel neuronal response was converted into the a feature representation and evaluated by a fixed, pretrained linear readout implemented on the CL1. The readout produced one of five steering classes, which was transmitted to the host PC and directly applied as the steering command for the subsequent game state. The corresponding reference steering class was recorded independently for comparison with the neuronal prediction. Readout training was performed offline, whereas all game-play evaluation was conducted in closed loop, such that each neuronal prediction directly influenced the subsequent game state and, consequently, the next input presented to the neuronal reservoir. Complete definitions of the game-state encoding, stimulation protocol, neuronal-response processing, readout model, and evaluation procedure are provided in Methods (Section 4.9).

#### 2.3.1 *Luigi Circuit* and *Mushroom Gorge* Actions, Tracking, and Completion Results

To evaluate whether the neuronal-reservoir predictions were sufficiently accurate for closed-loop game play, ten runs were performed on both *Luigi Circuit* and *Mushroom Gorge*, two of the 32 tracks in *Mario Kart Wii*. Performance was evaluated at two levels: agreement between the CL1-generated steering actions and the centerline-derived reference controller, and the resulting ability of the kart to remain near the track centerline and progress through the track. All runs used predictions generated by the same fixed readout trained offline using 4000 CL1 neuronal responses.

Across the ten *Luigi Circuit* runs, exact five-class agreement between the neuronal prediction and the reference steering action was 65.87%, while directional agreement was 74.89%, as shown in Fig. 8(a). Directional agreement groups steering actions according to their overall left, straight, or right direction and provides a measure of whether the reservoir selected the appropriate general steering direction even when the predicted steering magnitude differed from the reference. At the 5-Hz closed-loop control rate, the steering command was updated every 200 ms. Consequently, an isolated misclassification did not necessarily result in a sustained trajectory error, because subsequent neuronal predictions could modify the steering command and correct the kart’s trajectory.

**Fig. 8.**
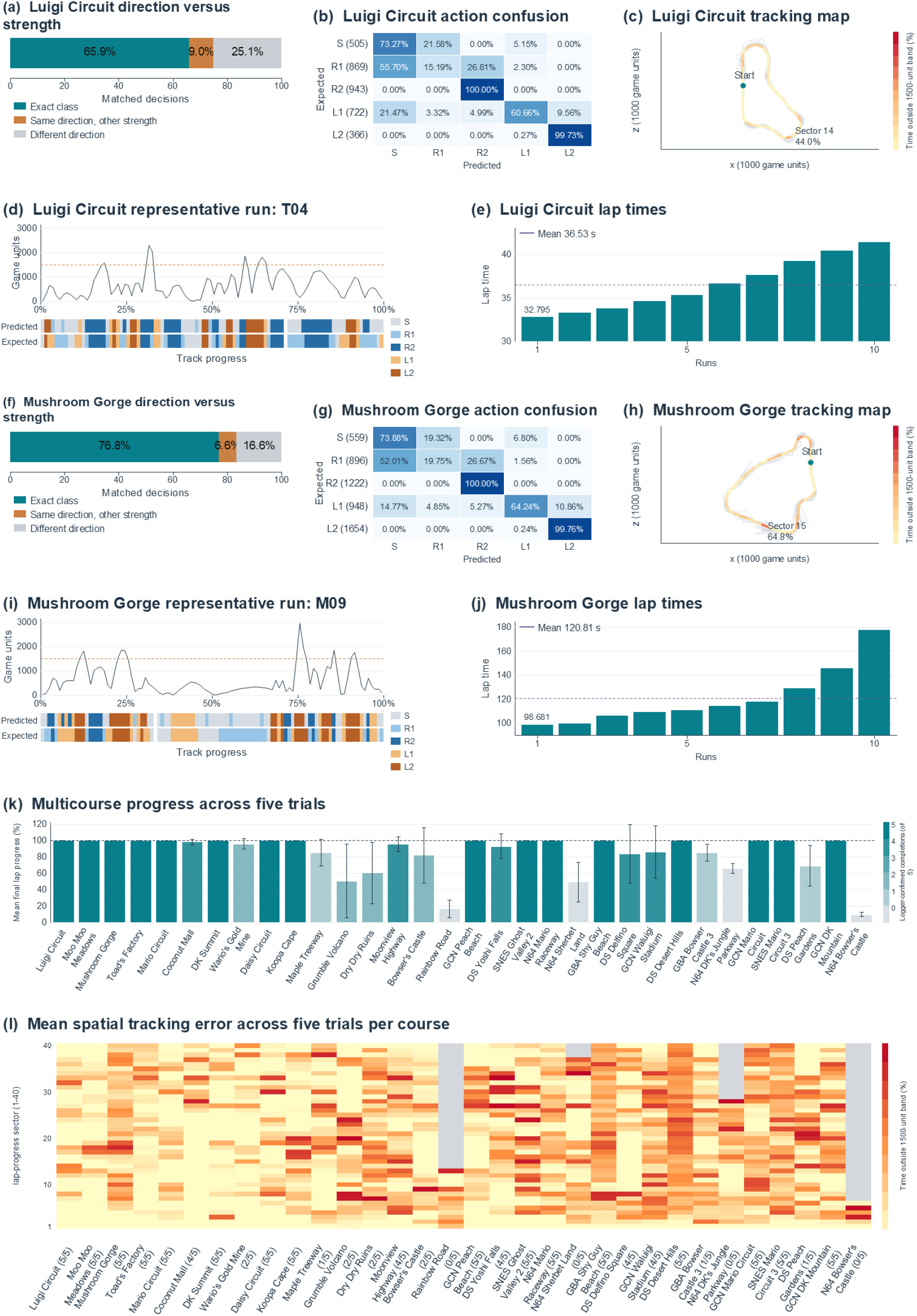
Closed-loop steering performance on tracks. (a) Luigi Circuit mean exact, same-direction/different-strength, and different-direction decision fractions. (b) *Luigi Circuit* five-class confusion matrix. (c) *Luigi Circuit* mean time outside the 1500-unit centerline band by track sector. (d) Tracking error and expected/predicted actions for run T04. (e) Lap time read at the visible transition from lap 1/3 to lap 2/3 for all ten *Luigi Circuit* runs. (f)–(j) The corresponding analyses for the ten *Mushroom Gorge* runs. (k) Mean final lap progress for exactly five trials per track. Error bars are one sample standard deviation (*n* = 5), and bar color gives the number of logger-confirmed completions out of five. (l) Pooled time-weighted centerline-band departures across 40 game-reported lap-progress sectors for the same five trials per track. Gray cells are sectors not observed in any of the five trials.

The pooled class-specific confusion matrix for Luigi Circuit is shown in Fig. 8(b). Correct predictions remained dominant across the ten runs, while most errors occurred between neighboring steering classes or between a turning action and the straight class. Thus, many classification errors represented differences in steering magnitude rather than a complete reversal of steering direction. For example, confusion between a soft and hard turn could cause the kart to respond more or less aggressively than the reference controller, whereas confusion between left- and right-directed actions would produce a substantially larger control error. The predominance of correct and directionally consistent predictions allowed the closed-loop system to maintain responsive steering despite an exact five-class agreement of 65.87

The resulting trajectory data demonstrate that this prediction performance was sufficient to support stable driving on *Luigi Circuit*. Across the ten runs, the kart spent an average of 86.47% of logged time within 1500 game units of the track centerline. Figure 8(c) shows the corresponding sector-level tracking performance. Sector 14 exhibited the largest deviation from the centerline, indicating that this region of the track produced the greatest tracking difficulty and coincided with a change in track geometry. These localized deviations did not prevent continued progression through the track, demonstrating that the closed-loop controller could recover from transient departures from the centerline.

An individual *Luigi Circuit* trajectory, run T04, is shown in Fig. 8(d). This run remained within the 1500-game-unit centerline band for 88.89% of the observation interval and achieved a video-verified first-lap time of 35.331 s. All ten *Luigi Circuit* runs successfully reached the visible transition from lap 1/3 to lap 2/3. First-lap times ranged from 32.795 to 41.441 s, with a mean of 36.533 s and a median of 36.008 s, as summarized in Fig. 8(e). The fastest observed first lap was completed in 32.795 s. These lap times were obtained directly from the frozen in-game display in the corresponding screen recordings and therefore exclude recorder lead-in, reset periods, and post-lap idling.

The same evaluation was performed on *Mushroom Gorge*, which introduced a more geometrically complex track and additional failure modes. Across the ten *Mushroom Gorge* runs, exact five-class agreement increased to 76.83%, while directional agreement reached 83.42%, as shown in Fig. 8(f). As with *Luigi Circuit*, all steering commands applied during these runs were produced directly by the fixed CL1 readout. The pooled class-specific confusion matrix in Fig. 8(g) shows strong agreement for both left- and right-directed steering actions, with the most prominent confusion occurring between soft-right and straight predictions. This pattern again indicates that a substantial fraction of the remaining errors altered steering magnitude rather than reversing the required steering direction.

Importantly, these agreement values quantify similarity between the CL1-generated actions and the centerline-derived reference controller rather than the optimality of either controller. The reference actions provide a consistent baseline against which neuronal predictions can be evaluated, but disagreement with the reference does not necessarily imply that the applied action was incapable of maintaining the kart on the track. Closed-loop performance must therefore also be considered in terms of the resulting trajectory and completion behavior.

Across the *Mushroom Gorge* runs, the kart spent an average of 74.00% of logged time within 1500 game units of the track centerline. This value was lower than the 86.47% observed on *Luigi Circuit*, consistent with the increased geometric and environmental complexity of *Mushroom Gorge*. The sector-level analysis in Fig. 8(h) identified Sector 15 as having the largest mean out-of-band fraction. Unlike *Luigi Circuit, Mushroom Gorge* contains regions in which the kart can leave the drivable surface, fall from the track, and be automatically repositioned at a designated reset location. These events introduce larger deviations from the centerline and additional recovery periods that are not present during ordinary steering corrections.

Run M09, shown in Fig. 8(i), remained within the 1500-game-unit centerline band for 78.99% of its logged duration and reached the finish with a logger-recorded control-to-completion time of 110.571 s. All ten *Mushroom Gorge* runs successfully reached the final-lap endpoint. Across the ten trials, logged control-to-finish times ranged from 98.681 to 177.405 s, with a mean of 120.809 s and a median of 112.444 s, as shown in Fig. 8(j). The broader completion-time distribution relative to the *Luigi Circuit* first-lap measurements is consistent with the more difficult track geometry, increased frequency of slowdowns and corrective steering, and the possibility of fall-and-reset events.

Taken together, these results demonstrate that the neuronal reservoir was capable of generating steering predictions that supported sustained closed-loop control across two distinct tracks. Although the CL1 readout did not reproduce the reference controller on every 200-ms control interval, most disagreements were either directionally consistent or could be corrected by subsequent predictions. The resulting trajectories show that interval-by-interval classification agreement alone does not fully characterize closed-loop performance: repeated neuronal predictions formed a continuous feedback process in which later actions could compensate for earlier errors. The successful progression of all *Luigi Circuit* runs through the first lap and completion of all ten *Mushroom Gorge* runs therefore demonstrates that the fixed neuronal readout generated sufficiently consistent control actions to support extended autonomous game play.

#### 2.3.2 Multi-Track Coverage and Communication Timing

Following the initial validation experiments on *Luigi Circuit* and *Mushroom Gorge*, the closed-loop neuronal controller was evaluated across all 32 tracks available in *Mario Kart Wii* to assess the generality of the control framework across a substantially broader range of track geometries and driving conditions. Importantly, the same overall neuronal-reservoir control framework was used across the full set of tracks, allowing performance to be evaluated beyond the tracks used for the initial detailed analysis. The resulting multi-track evaluation is summarized in Fig. 8(k,l).

A total of 160 trials were included in this analysis, consisting of exactly five trials for each of the 32 tracks. To prevent tracks with additional recorded attempts from receiving disproportionate weight, trials were selected strictly according to attempt index, with attempts 1–5 retained for every track. This produced an equally weighted evaluation in which each track contributed the same number of trials to the aggregate statistics.

Of the 160 selected trials, 111 resulted in logger-confirmed completion, corresponding to an overall trial-level completion rate of 69.38%. At least one successful completion was observed on 28 of the 32 tracks, while all five selected trials were completed successfully on 16 tracks. Thus, the neuronal controller achieved at least one successful run on 87.5% of the available tracks and achieved a 100% completion rate across the five selected trials on half of the tracks evaluated. These results demonstrate that the closed-loop controller was not limited to the two tracks examined in detail, but was capable of producing functional steering behavior across a broad range of previously unevaluated track layouts.

Because binary completion alone does not capture how far the kart progressed during unsuccessful trials, final track progress was also analyzed. Mean progress across the 32 equally weighted tracks was 85.27%, while the median track-level progress was 99.26%. Additionally, 20 of the 32 tracks exhibited a mean progress of at least 90% across their five selected trials. The high median relative to the mean indicates that many tracks were completed or nearly completed consistently, whereas a smaller number of more difficult tracks produced substantially lower progress and reduced the overall mean.

Figure 8(k) reports the mean and standard deviation of final progress for each track. Progress was normalized from the beginning of lap 1 to the lap 2 transition, providing a common measure that could be compared across tracks with different geometries and lengths. Under this definition, a progress value approaching 100% indicates that the kart reached or nearly reached the lap 2 transition during the evaluated interval. The standard deviation additionally characterizes run-to-run variability and identifies tracks for which successful progression was less consistent across the five trials.

Tracking behavior across the complete 32-track evaluation is shown in Fig. 8(l). For each track, observation time from the same five selected trials was pooled and divided into 40 sectors according to the game-reported lap-progress value. Within each sector, the figure reports the fraction of observed time for which the kart was more than 1500 game units from the corresponding track centerline. This representation provides a spatially normalized view of where tracking difficulty occurred and allows deviations to be compared across tracks despite differences in track geometry and length. A gray cell indicates that none of the five selected trials reached the corresponding sector; therefore, gray regions represent unavailable observations rather than zero tracking error. The multi-track results also highlight an important distinction between local steering accuracy and overall closed-loop driving capability. A neuronal prediction does not need to match the reference steering class at every control interval for the kart to continue progressing through the track. Because steering is repeatedly updated from the evolving game state, subsequent neuronal predictions can compensate for earlier deviations, provided that the accumulated error does not place the kart in an unrecoverable state. Consequently, the observation that 111 of 160 trials reached the evaluated completion criterion, and that 28 of 32 tracks produced at least one successful completion, provides a system-level measure of the ability of the neuronal reservoir to sustain control across diverse track configurations.

An additional constraint on closed-loop performance was the communication and processing delay between the game client and the CL1 system. Unlike a conventional controller operating entirely on the host computer, each neuronal control cycle required transmission of the encoded game state from the host PC to the remote CL1 system, stimulation of the neuronal culture, acquisition and processing of the resulting neuronal response, generation of a readout prediction, and transmission of the predicted steering action back to the game client. The measured round-trip time therefore captures the combined latency associated with client transmission, server-side processing, CL1 stimulation and neuronal-response acquisition, readout evaluation, return transmission, and final receipt by the game client.

Across the ten detailed *Luigi Circuit* runs, the median measured round-trip time was 408.27 ms. For the ten *Mushroom Gorge* runs, the corresponding median was 417.75 ms. These latencies are substantially longer than the nominal 200-ms interval associated with a 5-Hz steering-update rate and therefore represent an important practical limitation of the distributed closed-loop implementation. Nevertheless, successful sustained driving and track completion were observed despite this communication and neuronal-processing delay. The ability of the system to operate under these conditions indicates that the closed-loop steering task retained sufficient tolerance to delayed control updates for subsequent predictions to correct developing trajectory errors.

Taken together, the 32-track evaluation demonstrates that the neuronal-reservoir controller generalized beyond the individual tracks used for detailed characterization. The system produced successful or near-complete trajectories across the majority of *Mario Kart Wii* tracks while operating through a distributed control loop that incorporated neuronal stimulation, response acquisition, classification, and network communication. These results establish the multi-track experiment as a system-level demonstration of neuronal-reservoir control under varied track geometries and real-time communication constraints.

## 3 Discussion

Neuronal cultures are intrinsically dynamic and continuously evolving systems. Their spontaneous activity is spatially heterogeneous, their population state transitions between distinct dynamical regimes, and their functional organization changes over time. Computing with such a substrate must therefore be fundamentally different from computing with conventional hardware or static artificial reservoirs, where the underlying system is typically assumed to remain approximately invariant. Reliable biological computation consequently requires additional mechanisms for observing, stabilizing, adapting to, or controlling the state of the living network. However, strong external control, repeated retraining, or deliberate modification of the neuronal dynamics can make computation increasingly dependent on the imposed intervention and may suppress or obscure the intrinsic dynamical properties that make the biological substrate computationally interesting in the first place.

This study therefore takes a different approach: rather than attempting to force the neuronal network into a prescribed computational state, we perform computation on top of its unmodified native dynamics. We show that the naturally evolving culture already provides the essential ingredients of reservoir computation, including nonlinear transformation, transient memory, input-state separation, and rich state-dependent responses. Importantly, spontaneous network state is not treated merely as unwanted variability, but as part of the computational condition of the reservoir. The observation that different dynamical regimes support different levels of input separability further indicates that effective biological computation depends not only on how the network is stimulated, but also on *when* it is stimulated.

The broader scientific impact is a shift in how living neuronal systems can be engineered for computation. Rather than requiring the biological substrate to behave as stationary hardware, our results support a framework in which heterogeneity, metastability, and neural drift are explicitly measured and computationally accommodated. In this view, the objective is not to eliminate biological evolution, but to identify and exploit computationally favorable regions of an evolving dynamical system. This provides a foundation for living computing architectures in which computation is continuously aligned with the native state of the biological substrate rather than imposed through persistent modification of it.

## 4 Methods

### 4.1 Neuronal Culture and MEA Experimental Setup

Electrophysiological experiments were performed using the CL1 neurocomputational platform (Cortical Labs, Melbourne, Australia), which integrates living neuronal cultures, multielectrode-array (MEA) recording and stimulation, environmental regulation, and programmable closed-loop experimental control within a single system [23]. The neuronal preparation consisted of approximately 2 × 10^5^ human iPSC-derived NGN2 cortical neurons co-cultured with approximately 4 × 10^4^ primary human cortical astrocytes. This co-culture formed the recurrent neuronal network used as the physical reservoir throughout the study.

Neuronal activity was interfaced through the standard CL1 MEA. The platform provides 59 simultaneously addressable measurement electrodes together with a reference electrode; electrode pads are 30 *µ*m in diameter with a 200 *µ*m pitch over an active area of approximately 1.4 × 1.4 mm^2^. Extracellular potentials were sampled simultaneously at 25 kHz with a measurement bandwidth of 200–10,000 Hz. The acquisition circuitry provides a ± 5 mV input range with 0.195 *µ*V resolution. Raw electrophysiological signals, detected neuronal events, and delivered stimulation events were stored in HDF5 format for subsequent analysis [23].

Electrical perturbations were delivered through the CL1 programmable constant-current stimulation interface. The system supports independently configurable monophasic, biphasic, and triphasic stimulation waveforms across the stimulation-capable electrodes, with currents up to ± 12.75 *µ*A, 50 nA current resolution, and 20 *µ*s pulse-width resolution [23]. Charge-balanced biphasic stimulation was used in the present study, with stimulation location, current amplitude, pulse width, frequency, and pulse number varied according to the corresponding experimental protocol described below.

The neuronal culture was maintained on the integrated CL1 life-support stage during recording and stimulation. The MEA surface was maintained at 37°C and the atmospheric environment was regulated at a nominal CO_2_ concentration of 5%. Oxygen was not actively controlled and remained approximately at atmospheric concentration during experimentation. The CL1 platform provides integrated temperature and gas regulation for long-term neuronal-culture operation. Experimental acquisition, stimulation, and closed-loop control were implemented using Python through the device-hosted Jupyter programming environment and CL1 application programming interface, which provides real-time access to electrophysiological recordings and programmable stimulation [23].

### 4.2 Spontaneous Network Dynamics Analysis

Spontaneous neuronal activity was analyzed at the population level to characterize both the geometry of the evolving reservoir state and the functional interaction structure among recording sites. Raw extracellular recordings were first converted to channel-wise spike trains using the adaptive detection procedure described in Section 4.3. Unless otherwise stated, negative-polarity events were detected using a threshold factor of *k* = 5.0 and a refractory interval of 1 ms. The resulting spike times were subsequently transformed into temporally resolved firing-rate representations that formed the basis of both the population state-space and functional-connectivity analyses.

#### 4.2.1 Population State-Space Analysis

To obtain a low-dimensional representation of the spontaneous neuronal population dynamics, detected spikes were converted into a time-resolved firing-rate matrix. PCA and related dimensionality-reduction methods are widely used to summarize coordinated structure in high-dimensional neural population recordings [24], including spontaneous activity in cultured neuronal networks [25]. The recording was divided into non-overlapping temporal bins of duration

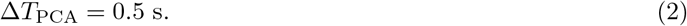

For recording channel *i* and temporal bin *b*, the firing rate was defined as

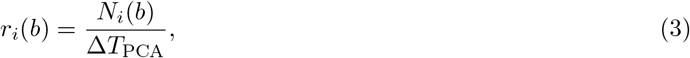

where *N*_*i*_(*b*) denotes the number of detected spikes on channel *i* within bin *b*. The instantaneous population state associated with that bin was therefore represented by

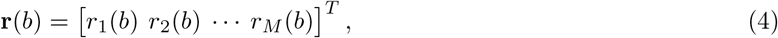

where *M* denotes the number of indexed recording channels included in the analysis. Stacking the population vectors over all *B* temporal bins produced the firing-rate matrix

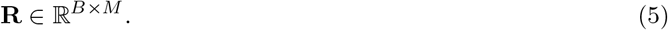

Because spontaneous firing rates differed substantially across recording locations, each channel was standardized independently before dimensionality reduction. For channel *i*, the standardized firing rate was

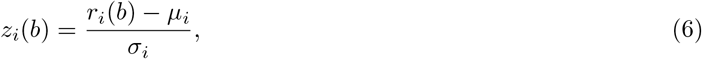

where *µ*_*i*_ and *σ*_*i*_ are the mean and standard deviation, respectively, of the firing-rate series of channel *i* across temporal bins. The resulting standardized population matrix **Z** therefore assigns comparable statistical scale to all recording dimensions while preserving their temporal covariation.

Principal component analysis (PCA) was then applied to **Z** to identify orthogonal directions accounting for the dominant variance in spontaneous population activity [24]. Let **W** denote the matrix containing the eigenvectors corresponding to the leading principal components. The low-dimensional population state at bin *b* was represented as

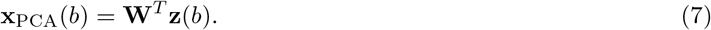

Two- and three-dimensional projections were generated using the leading two and three principal components, respectively. Each projected point therefore represents the population firing state during one 0.5-s interval, and points were indexed by recording time to visualize the temporal evolution of the spontaneous reservoir trajectory. The fraction of total variance explained by each retained principal component was calculated from the corresponding PCA eigenvalue spectrum.

PCA was fitted independently to each spontaneous recording. Consequently, comparisons between recordings or experimental days were interpreted in terms of changes in state-space geometry, dispersion, and variance structure rather than as direct correspondence between absolute principal-component coordinates.

#### 4.2.2 Functional Connectivity Analysis

Functional connectivity was estimated from correlations between the same temporally binned firing-rate signals used for the population-state analysis. Correlation-based inference from MEA spike activity is an established approach for estimating functional interactions in cultured neuronal networks [25, 26]. For each pair of recording channels *i* and *j*, the Pearson correlation coefficient was calculated across the *B* firing-rate bins,

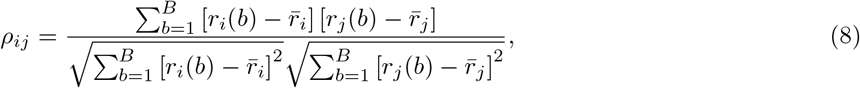

where 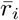 and 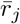denote the mean firing rates of channels *i* and *j*, respectively. The resulting matrix

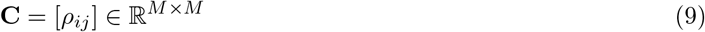

provides a symmetric measure of statistical coupling between electrode firing-rate fluctuations. Undefined correlations resulting from channels with zero temporal variance were assigned a value of zero.

An undirected functional network was constructed from **C**. Recording channels were represented as graph nodes, and an edge between channels *i* and *j* was retained when the magnitude of their firing-rate correlation exceeded the predefined threshold

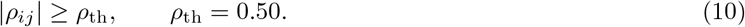

Thus, both sufficiently strong positive and negative correlations contributed to the functional-connectivity graph. The absolute correlation magnitude,

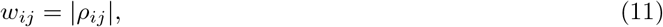

was used as the edge strength for visualization, such that more strongly correlated channel pairs were represented by thicker connections. Nodes were positioned according to the indexed MEA grid layout to preserve the approximate spatial organization of the recording array.

Functional-connectivity matrices and networks were calculated independently for each spontaneous recording. Changes between recording sessions were therefore interpreted as changes in the statistical interaction structure of the neuronal population, rather than as direct evidence of anatomical synaptic connectivity [26], providing a complementary measure of network reorganization alongside the PCA-based population state-space analysis.

### 4.3 Adaptive spike detection

Extracellular recordings obtained from different MEA electrodes exhibit substantial differences in baseline voltage, background fluctuation amplitude, and effective signal-to-noise ratio. A single absolute threshold would therefore impose different effective detection sensitivities across recording sites. To obtain an approximately channel-invariant event representation, spike detection was performed independently for each electrode using a robust median-absolute-deviation (MAD)-based threshold, following established robust extracellular spike-detection principles [27–29]. The complete processing sequence consisted of channel-wise background estimation, adaptive thresholding, threshold-crossing detection, refractory-period enforcement, and temporal integration of the detected events into reservoir-state vectors.

Throughout this section, *i ∈* {1, …, *M*} denotes the recording-channel index, *n ∈* { 0, …, *N* − 1 } denotes the discrete sample index, *M* is the number of recording channels, *N* is the number of samples in the analyzed recording segment, and *f*_*s*_ is the sampling frequency in Hz. The physical time associated with sample *n* is *t*_*n*_ = *n/f*_*s*_.

#### 4.3.1 Channel-wise robust background estimation

For recording channel *i*, let *V*_*i*_[*n*] denote the discrete extracellular voltage signal. The channel-specific baseline *m*_*i*_ was estimated using the median, *m*_*i*_ = median_*n*_ [*V*_*i*_[*n*]].

The corresponding median absolute deviation was

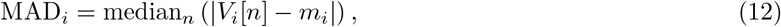

and was converted to a Gaussian-equivalent robust noise estimate,

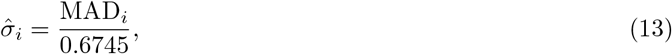

where 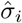 represents the estimated background-noise scale of channel *i*. The factor 0.6745 makes the MAD estimator consistent with the standard deviation for a Gaussian distribution. Median-based estimators were used because they are less sensitive than variance-based estimators to large transient extracellular events and occasional outliers [27–29].

For the negative-polarity extracellular events analyzed here, the adaptive detection threshold for channel *i* was defined as

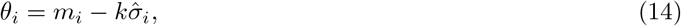

where *θ*_*i*_ is the channel-specific voltage threshold and *k* is the dimensionless threshold factor.

After median centering, such that 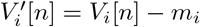, the corresponding threshold becomes

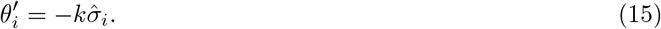

Thus, event detection was based on the magnitude of a voltage excursion relative to the local statistical background of each electrode rather than on a common absolute voltage level. For two electrodes *i* and *j* having different background-noise scales,

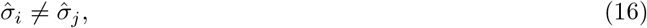

an absolute voltage excursion with magnitude |*A*| corresponds to the normalized channel-specific amplitude

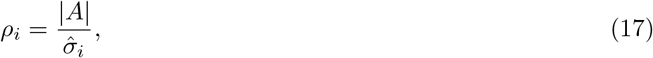

where *ρ*_*i*_ is dimensionless. This dependence motivates channel-wise normalization of the detection threshold.

#### 4.3.2 Threshold-crossing detection and refractory-period enforcement

For negative-polarity detection, a candidate event occurred at sample *n* when the signal crossed the adaptive threshold from above,

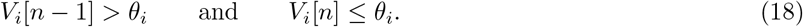

The candidate-crossing set for channel *i* was therefore

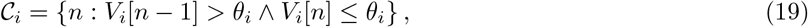

where *C*_*i*_ contains the sample indices of all candidate threshold crossings on channel *i*.

Because a single extracellular waveform can generate closely spaced threshold crossings, a refractory interval *T*_ref_ was imposed.

The corresponding minimum number of samples between accepted events was

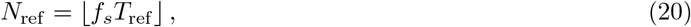

where *N*_ref_ is the refractory interval expressed in samples.

If *n*_*i,j*_ denotes the sample index of the *j*th accepted event on channel *i*, consecutive accepted events were required to satisfy

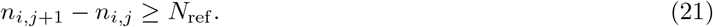

Defining the corresponding event time as

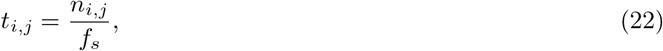

the equivalent continuous-time constraint is

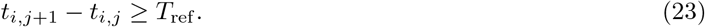

The resulting spike train for electrode *i* was represented as the point process

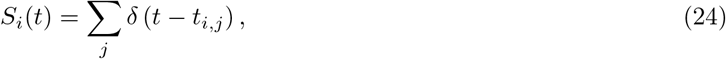

where *j* indexes accepted events and *δ*(·) denotes the Dirac delta function.

#### 4.3.3 Response-window integration and reservoir-state construction

The detected events were integrated over finite post-stimulation response windows. For a response window of duration *W* beginning at time *t*_0_, the event count on channel *i* was

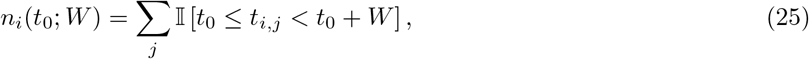

where *W* is the response-window duration and I[·] is the indicator function, equal to one when its argument is true and zero otherwise.

The population reservoir state was then constructed as

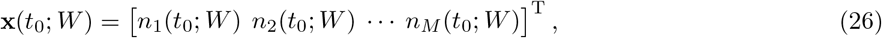

where **x**(*t*_0_; *W* ) *∈* ℝ^*M*^ is the measured population reservoir-state vector. Accordingly, the measurement pipeline can be summarized as

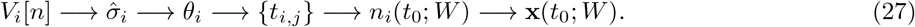

The threshold factor *k* determines which voltage excursions enter the detected-event set, whereas *W* determines the temporal scale over which these events are integrated to form the measured reservoir state.

#### 4.3.4 Synthetic spike-injection validation

To quantify detector sensitivity under controlled conditions, synthetic extracellular spike waveforms of known timing and amplitude were superimposed on empirically recorded channel backgrounds, consistent with the use of simulated or semi-simulated signals for controlled spike-detector validation [27, 28]. This validation does not provide biological spike ground truth; rather, it tests whether the same known waveform can be detected consistently across channels having different background-noise levels.

A validation subset *V* containing

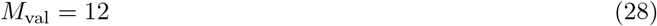

recording channels was selected approximately uniformly across the empirical distribution of 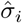,using noise quantiles spanning the low- to high-noise range. Thus, |*V* | = *M*_val_.

For each selected channel, an 8-s background segment was extracted from the median-centered voltage trace,

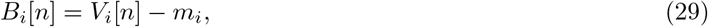

where *B*_*i*_[*n*] denotes the empirical background used for synthetic validation.

To prevent rare extreme excursions from dominating the validation, background samples were clipped according to

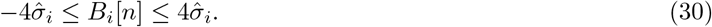

Each synthetic validation trace contained

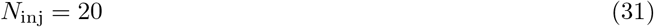

injected events, and the validation procedure was repeated

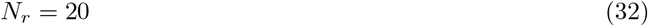

times for every tested channel and normalized amplitude level, where *r∈* { 1, …, *N*_*r*_} denotes the validation-repeat index.

#### 4.3.5 Synthetic extracellular spike waveform

The injected waveform was a normalized biphasic extracellular-spike template with a dominant negative phase followed by a smaller delayed positive phase. Prior to normalization, the waveform was defined as

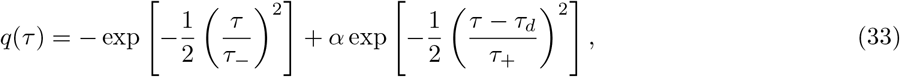

where *τ* is time relative to the synthetic-spike center and

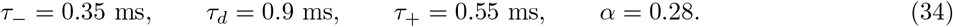

Here, *τ*_*−*_ controls the width of the dominant negative phase, *τ*_*d*_ is the delay of the positive phase, *τ*_+_ controls its width, and *α* specifies its relative amplitude.

The template was evaluated over

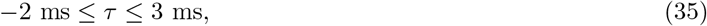

giving a waveform-support duration

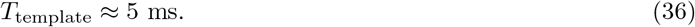

The template was normalized to unit absolute peak amplitude,

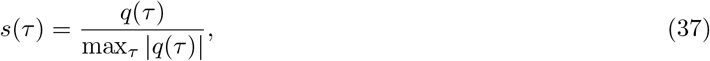

where *s*(*τ* ) is the normalized continuous-time spike template. Sampling *s*(*τ* ) at the recording frequency *f*_*s*_ produced the discrete template *s*[*ℓ*], where *ℓ* denotes the template-sample index.

#### 4.3.6 Injected amplitude and signal-to-background scaling

Synthetic spikes were evaluated over the dimensionless normalized amplitude levels

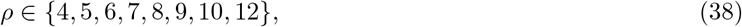

where *ρ* denotes injected spike amplitude relative to the local MAD-derived noise scale. For channel *i*, the absolute injected waveform amplitude was

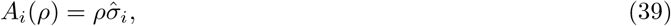

where *A*_*i*_(*ρ*) has the same voltage units as *V*_*i*_[*n*].

The synthetic trace was therefore

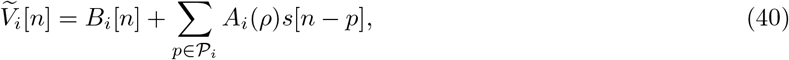

where 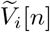 is the synthetic validation trace and _*i*_ denotes the set of known injection sample indices for the corresponding validation trace.

Thus, *ρ* = 7 denotes an injected spike whose peak amplitude is seven times the local MAD-derived background scale of that channel. This quantity should not be confused with the electrical stimulation current delivered to the neuronal culture.

Injection locations were selected away from the boundaries of the synthetic trace and were separated by at least

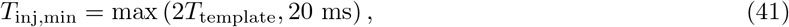

where *T*_inj,min_ denotes the minimum temporal separation between synthetic injections. Since *T*_template_ ≈ 5 ms, the resulting minimum separation was 20 ms.

#### 4.3.7 Adaptive and global detector comparison

The adaptive detector used the channel-specific noise scale 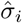 for each electrode. For comparison, the global detector used a single noise scale defined as the median across the *M*_val_ validation channels,

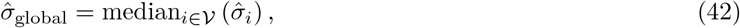

where 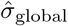 is the common background-noise estimate used by the global detector.

After median centering, the corresponding global negative threshold was

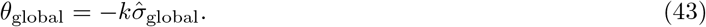

The adaptive and global methods therefore differed only in whether the detection scale was electrode specific or shared across validation channels.

#### 4.3.8 Injected-spike detection rate

Because the injection times were known exactly, each synthetic event could be matched against the detector output. A detected event was considered a successful match when it occurred within a temporal tolerance

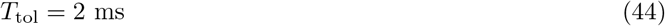

of the corresponding injected event.

For validation channel *i*, normalized amplitude *ρ*, and repeat *r*, the injected-spike detection rate was defined as

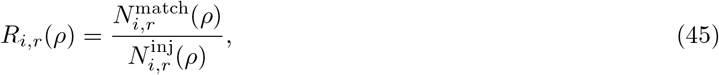

where 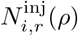 is the number of synthetically injected spikes and 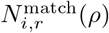 is the number of injected spikes for which a detected event occurred within ± *T*_tol_.

For every validation trace,

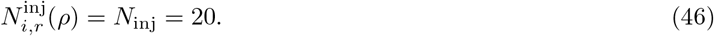

For each channel, the repeated detection-rate estimates were first averaged,

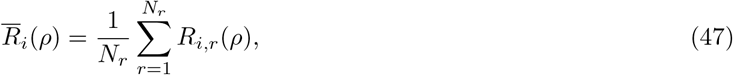

where 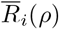 denotes the repeat-averaged detection rate for channel *i* at normalized amplitude *ρ*.

This averaging was performed before computing across-channel variability so that repeat-to-repeat stochasticity was not conflated with electrode-to-electrode heterogeneity.

The mean detection rate across the validation channels was then

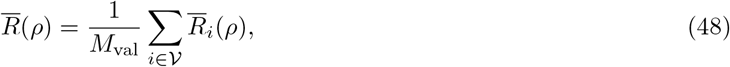

where 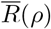 represents the population-averaged injected-spike detection rate.

#### 4.3.9 Across-channel detection variability

To determine whether detector performance depended on recording location, the sample variance of the channel-averaged detection rates was calculated at each normalized injected amplitude,

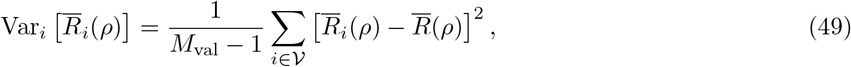

where Var_*i*_ [.] denotes variance across validation channels. The corresponding across-channel standard deviation was

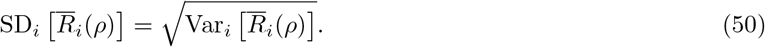

Relative channel-to-channel variability was quantified using the coefficient of variation,

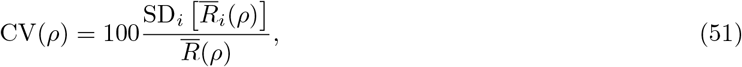

where CV(*ρ*) is expressed as a percentage. The coefficient of variation was treated as undefined when 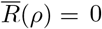, rather than introducing an artificial regularization term.

To directly compare the global and adaptive thresholding approaches, the across-channel variance ratio was defined as

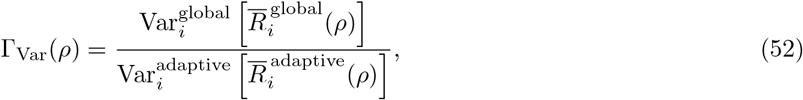

where the superscripts global and adaptive identify the corresponding detection methods. Values

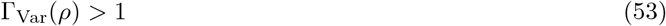

indicate greater electrode-dependent variability under the global detector. An analogous ratio was defined for the coefficient of variation,

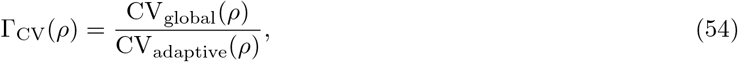

where Γ_CV_(*ρ*) *>* 1 indicates greater relative across-channel variability under global thresholding.

These controlled validation metrics distinguish two aspects of detector performance: 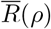 quantifies average sensitivity to known injected events, whereas Var_*i*_ 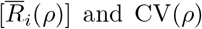 and CV(*ρ*) quantify the absolute and relative dependence, respectively, of that sensitivity on electrode-specific background statistics.

#### 4.3.10 Interpretation for reservoir-state construction

The principal objective of adaptive detection was not simply to maximize the total number of detected events, but to reduce systematic differences in effective detection sensitivity across electrodes. Because each recording channel contributes one dimension to the reservoir-state vector in Eq. 26, electrode-dependent detection bias can deform the measured geometry of the population state before any computational analysis is performed. Channel-wise MAD normalization therefore acts as a measurement normalization layer that allows differences among the dimensions of **x**(*t*_0_; *W* ) to more faithfully reflect neuronal activity rather than differences in local voltage scale or background noise.

### 4.4 State Awareness

To account for the spontaneous non-stationarity of the neuronal culture, the instantaneous network condition was estimated continuously from population spiking activity and classified into three operational dynamical regimes: *random/sparse, near-chaotic*, and *synchronized*. Cultured cortical networks are known to exhibit spontaneous population bursts, synchronized network events, and preparation-dependent changes in collective activity [30, 31]. State detection was based jointly on the temporal concentration of population spikes, the fraction of participating electrodes, and the total population spike count. Importantly, the decision boundaries were not fixed a priori; they were calibrated from the spontaneous activity of each preparation before state-aware stimulation.

#### 4.4.1 Selection of state-detection electrodes

Before state-aware stimulation, spontaneous activity was recorded for

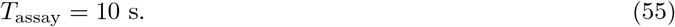

Electrodes excluded from stimulation or recording analysis were first removed from the candidate set. For each remaining electrode *i*, the number of spontaneous spikes observed during the assay was counted,

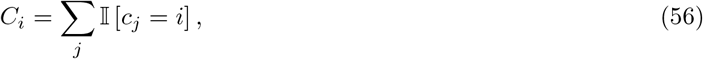

where *c*_*j*_ denotes the electrode associated with spike *j* and I[·] is the indicator function. The electrodes were ranked according to *C*_*i*_, and the

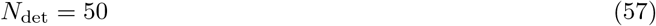

most active electrodes were retained as the state-detection population,

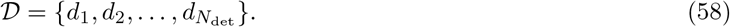

This procedure restricted state estimation to recording sites exhibiting measurable spontaneous activity while retaining a distributed population-level representation of the culture.

#### 4.4.2 Temporal population representation

State descriptors were calculated over a sliding observation window of duration

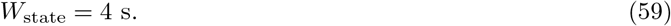

Within each window, spike times from the detection-electrode population were divided into bins of width

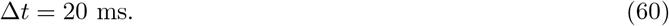

For a state-estimation time *t*, the analysis interval was therefore

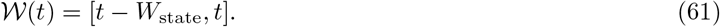

With *W*_state_ = 4 s and Δ*t* = 20 ms, each state estimate contained approximately

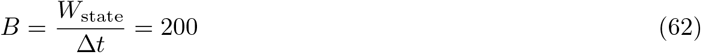

population-spike bins.

Let *N*_*b*_(*t*) denote the number of spikes from all detection electrodes occurring in temporal bin *b* within *W* (*t*).

The resulting population-count vector was

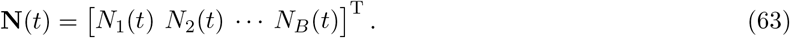

Three complementary population descriptors were then extracted from this window.

#### 4.4.3 Population spike count

The total number of detected spikes within the current state window was defined as

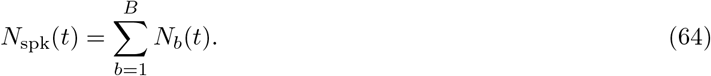

This quantity characterizes the overall level of population activity during the preceding 4 s.

#### 4.4.4 Active-electrode fraction

The spatial participation of the network was quantified as the fraction of detection electrodes contributing at least one spike during the current window. Let

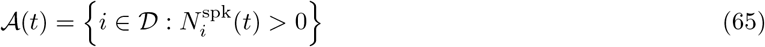

denote the subset of active detection electrodes, where 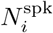 (*t*) is the spike count of electrode *i* within W(*t*). The active fraction was then

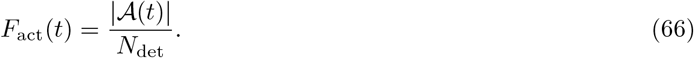

Thus,

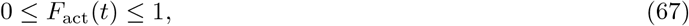

with values approaching unity indicating participation of most state-detection electrodes.

#### 4.4.5 Population synchronization index

Temporal coordination of population firing was quantified from the variability of the binned population spike counts, motivated by the strong temporal concentration and broad neuronal recruitment characteristic of synchronized network events in cultured cortical networks [30]. The synchronization index was defined as the coefficient of variation of **N**(*t*),

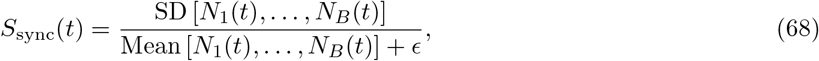

where

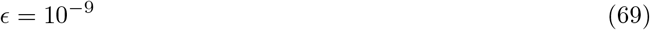

prevents division by zero.

The interpretation of *S*_sync_(*t*) follows directly from the temporal distribution of population events. If spikes are distributed relatively uniformly across the 20-ms bins, the variance of the population-count sequence is comparatively small and *S*_sync_ remains low. In contrast, strongly coordinated population bursts concentrate large numbers of spikes into a restricted subset of bins, increasing the temporal variance of **N**(*t*) and consequently producing a larger synchronization index.

It should therefore be noted that *S*_sync_ is a dimensionless population-event concentration metric rather than a pairwise spike-time synchrony coefficient.

#### 4.4.5 Preparation-specific calibration of state boundaries

Because spontaneous activity levels differed among neuronal preparations, the state-classification thresholds were estimated adaptively from the initial 10-s spontaneous assay rather than being imposed as fixed numerical constants. Preparation-adaptive detection has previously been used to account for substantial culture-to-culture variation in bursting and network-burst statistics [31].

During calibration, the same 4-s analysis window was evaluated at increments of

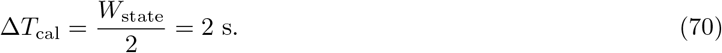

For each calibration window *q*, the three descriptors

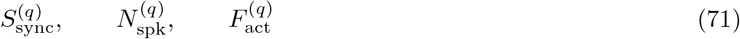

were calculated.

Four preparation-specific state thresholds were then obtained from the empirical distributions of these quantities. The lower synchronization boundary was defined as the 25th percentile of the synchronization-index distribution,

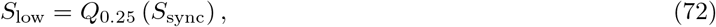

whereas the upper synchronization boundary was defined as its 75th percentile,

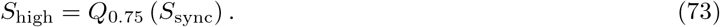

The low-activity spike-count threshold was similarly defined as

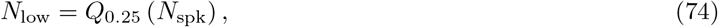

and the high-participation threshold was

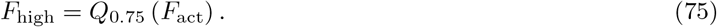

Here, *Q*_*p*_(*X*) denotes the empirical *p*th quantile of variable *X*. This percentile-based calibration makes the operational state boundaries preparation dependent and therefore responsive to differences in the intrinsic firing statistics of individual cultures.

#### 4.4.7 Operational state classification

At each state-estimation time *t*, the network was assigned to one of three dynamical regimes according to the joint values of *S*_sync_(*t*), *N*_spk_(*t*), and *F*_act_(*t*).

The state classifier can be written as

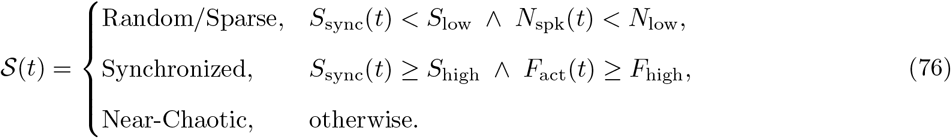

If no spikes were present within the current 4-s buffer, the network was assigned the auxiliary state

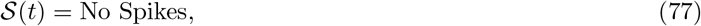

with

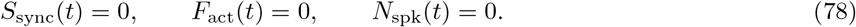

The *random/sparse* condition therefore required both low temporal concentration of population events and a low overall spike count. The *synchronized* condition required both a high synchronization index and broad spatial recruitment of the detection-electrode population. States not satisfying either extreme criterion were operationally assigned to the intermediate *near-chaotic* regime.

The term *near-chaotic* is used here as an operational state label defined by Eq. 76; the classifier itself does not constitute a direct estimate of a Lyapunov exponent or other formal chaos invariant. The terminology is motivated by theoretical and experimental work linking computationally rich recurrent dynamics to regimes near transitions between strongly ordered and irregular/critical activity [14, 32].

#### 4.4.8 Real-time state tracking

During state-aware experiments, spikes originating from the selected detection electrodes were continuously maintained in a rolling buffer,

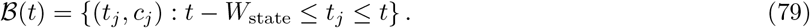

Events older than 4 s were removed continuously, ensuring that each state estimate reflected only the recent spontaneous history of the network.

Although acquisition proceeded at a higher internal loop rate, state classification was updated at

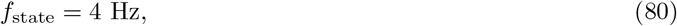

corresponding to one state estimate every approximately

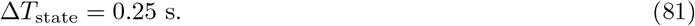

For every update, the instantaneous values

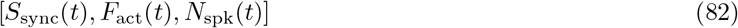

and the resulting state *S* (*t*) were stored in the state log. These measurements generated the temporal state trajectory used for subsequent state-occupancy and dwell-time analyses.

#### 4.4.9 State-conditioned stimulation

For state-aware stimulation experiments, the target operating regime was defined as

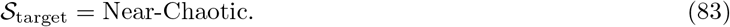

A stimulation event was permitted only when

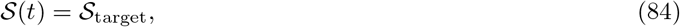

and when the required minimum interval from the previous stimulation had elapsed. The minimum inter-sample stimulation gap was

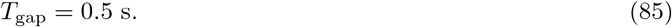

Consequently, the state-aware stimulation rule was

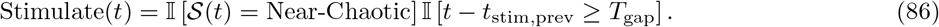

Immediately before each stimulation, the detected state and its underlying population descriptors were stored together with the stimulation record,

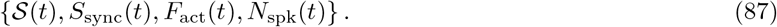

This provided an explicit record of the dynamical condition under which every external perturbation was applied and enabled subsequent comparison of stimulation-evoked reservoir responses as a function of the pre-stimulation network state.

### 4.5 Input Channel selection

Prior to the spatio-parametric stimulation experiments, candidate stimulation electrodes were screened to identify sites that reliably recruited distributed neuronal responses while avoiding the selection of multiple electrodes producing highly redundant population-response patterns. Site-specific electrical perturbation and stimulation-evoked population responses are well established in cultured neuronal networks interfaced through MEAs [33, 34]. Screening was performed independently of the subsequent state-conditioned stimulation protocol.

#### 4.5.1 Candidate electrode set

For stimulation screening, the acquired data structure was indexed over *N* = 64 channel positions. Five indices that were unavailable for stimulation were excluded,

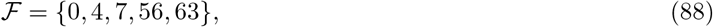

giving the candidate stimulation set

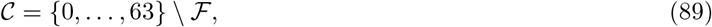

with

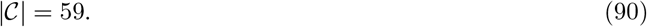

Each candidate electrode *c ∈ C* was evaluated independently using the same stimulation waveform and number of repeated trials.

#### 4.5.2 Screening stimulation protocol

For every candidate stimulation electrode, a charge-balanced biphasic pulse train was applied using a current amplitude of

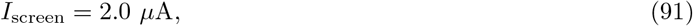

a phase duration of

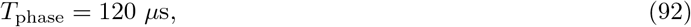

and

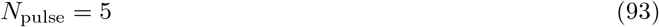

pulses delivered at

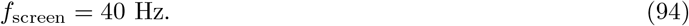

The biphasic stimulation waveform for each pulse can be represented as

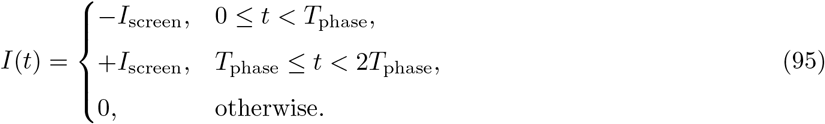

Each electrode was stimulated

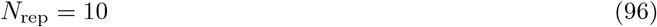

times, with a 1-s observation interval following each screening stimulation. Charge-balanced patterned electrical stimulation of cultured cortical networks has been widely used to evoke and characterize distributed network responses [33, 34].

#### 4.5.3 Evoked population-response vector

For each screening trial, stimulation-evoked spikes were counted during the post-stimulation interval

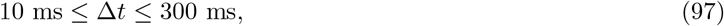

where

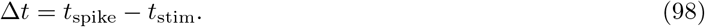

Spikes detected on the stimulating electrode itself were excluded from the screening response, as were electrodes outside the candidate set.

For stimulation electrode *c* and repetition *r*, the evoked population-response vector was

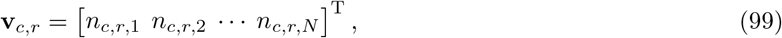

where *n*_*c,r,i*_ denotes the number of spikes detected on electrode *i* within the evoked-response interval after stimulating electrode *c* during repeat *r*. By construction,

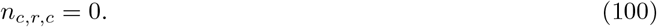

The mean spatial response map of stimulation electrode *c* was then calculated as

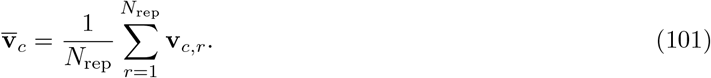

This vector characterizes the average spatial pattern of neuronal recruitment generated by stimulation at electrode *c*.

#### 4.5.4 Evoked-response magnitude

The total evoked response during repetition *r* was defined as the sum of the post-stimulation spike counts across recording electrodes,

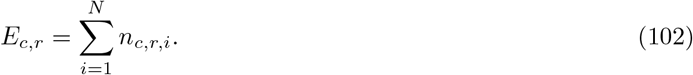

The mean evoked-response magnitude was

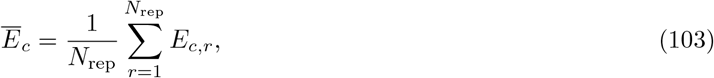

and its trial-to-trial standard deviation was

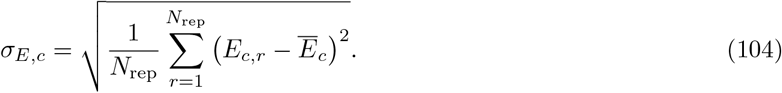

The implementation uses the population standard deviation for this quantity.

#### 4.5.5 Response reliability

A screening trial was considered responsive if at least one evoked spike was detected across the non-stimulating candidate-electrode population. The response reliability of electrode *c* was therefore

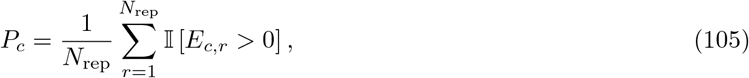

where I [·] denotes the indicator function.

Thus,

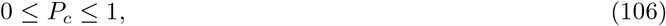

with *P*_*c*_ = 1 indicating that stimulation of electrode *c* evoked measurable activity on every screening repetition.

#### 4.5.6 Spatial recruitment

The spatial spread of the response was quantified from the mean response map in Eq. 101. The number of recruited electrodes was

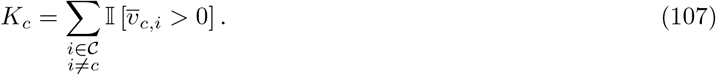

The corresponding active fraction was normalized by the number of available non-stimulating candidate electrodes,

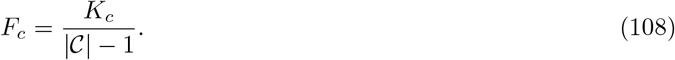

For the present electrode configuration,

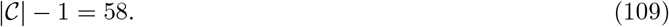

Therefore, *F*_*c*_ quantifies the fraction of the available electrode population recruited by stimulation of electrode *c*.

#### 4.5.7 Composite screening score

Each stimulation electrode was assigned a composite score combining response magnitude, repeatability, and spatial recruitment,

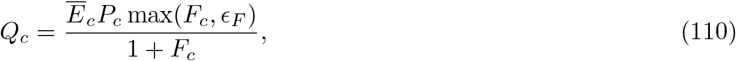

where

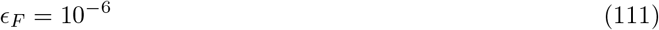

prevents an exactly zero spatial fraction from producing numerical degeneracy. The factor

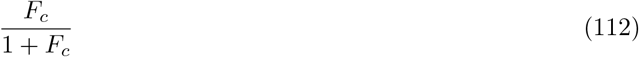

increases monotonically with spatial recruitment while sublinearly weighting increasingly widespread responses. Consequently, a high screening score requires three simultaneous properties: a large evoked population response *Ē*_*c*_, reliable recruitment across repeated stimulations *P*_*c*_, and distributed activation of the recording population *F*_*c*_.

Candidate electrodes were ranked in descending order of *Q*_*c*_.

#### 4.5.8 Response-diversity constraint

Ranking by response strength alone can select multiple stimulation sites that evoke nearly identical population patterns. To preserve spatial diversity among the selected inputs, the mean response maps of candidate electrodes were therefore compared using Pearson correlation.

For two stimulation electrodes *c* and *j*, the response-map correlation was

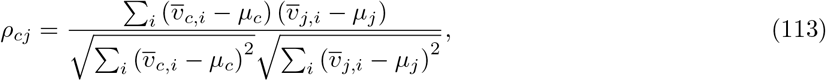

where

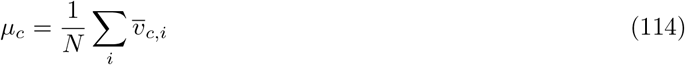

is the mean component of response map 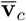.

If either response map had zero variance, its correlation with the comparison map was set to zero in the implementation.

Electrodes were considered response redundant when

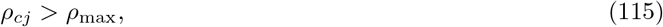

with

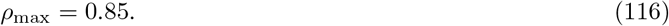

Beginning with the highest-scoring stimulation electrode, subsequent candidates were accepted only if their response-map correlation remained below or equal to 0.85 with every electrode already selected,

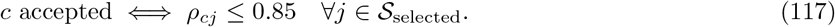

This greedy selection procedure preferentially retained electrodes that combined high screening scores with distinct spatial recruitment patterns.

#### 4.5.9 Final stimulation-electrode set

For the 10-class experiments, one stimulation electrode was assigned to each input class,

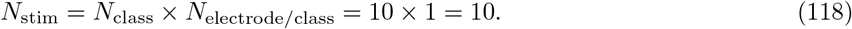

Screened electrodes were therefore added sequentially until

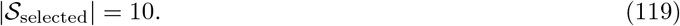

If the correlation-based diversity criterion yielded fewer than the required number of stimulation electrodes, the remaining positions were filled according to descending screening score irrespective of the correlation constraint.

For each screened electrode, the analysis retained the mean evoked response, trial-to-trial standard deviation, response reliability, spatial spread, active fraction, and composite screening score. These quantities were subsequently used to assess screening convergence and to define the stimulation sites employed in the spatio-parametric reservoir experiments.

### 4.6 Nonlinearity test

Nonlinear transformation is a central requirement of reservoir computation because it allows the recurrent substrate to map inputs into representations not available through a purely linear transformation [4, 6, 16]. For each recording channel *i*, the response expected under linear superposition was defined as

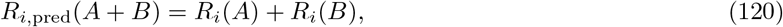

where *R*_*i*_(*A*) and *R*_*i*_(*B*) denote the mean evoked spike counts for the individual stimulation conditions. The channel-wise nonlinearity index was then calculated as

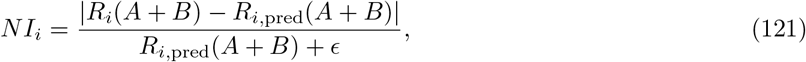

where *ϵ* = 10^*−*9^ prevents division by zero. A value of *NI*_*i*_ = 0 indicates exact linear superposition, whereas larger values indicate progressively stronger deviation from linearity.

### 4.7 Reservoir State Separability

Reservoir-state separability was evaluated by determining whether distinct spatio-parametric stimulation conditions were transformed by the neuronal culture into distinguishable multichannel population responses. The separation property is a core requirement of liquid-state and reservoir-computing systems and has been demonstrated experimentally in cultured neuronal networks [5–7]. Each input class was defined jointly by the spatial location of stimulation and the temporal structure of the applied electrical perturbation. The resulting analysis therefore characterized the input–state transformation

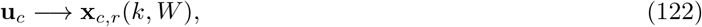

where **u**_*c*_ denotes stimulation class *c*, **x**_*c,r*_ is the measured reservoir state generated during repetition *r, k* is the spike-detection threshold factor, and *W* is the post-stimulation response-window duration.

### 4.7.1 Spatio-parametric input construction

Let

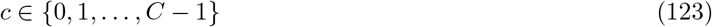

denote the stimulation-class index, where *C* is the number of distinct input classes. Following the stimulation-electrode screening procedure described in Section 4.5, one or more selected stimulation electrodes were assigned to each input class. The spatial component of class *c* was therefore represented by the corresponding stimulation-electrode set

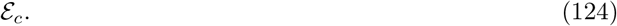

Each input class was additionally assigned a class-specific temporal stimulation configuration comprising stimulation frequency, number of pulses per burst, and pulse phase width. These parameters were distributed across the configured experimental ranges such that

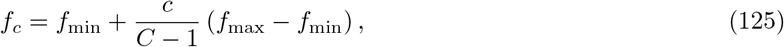

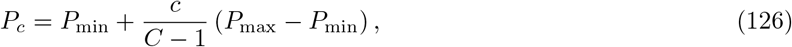

and

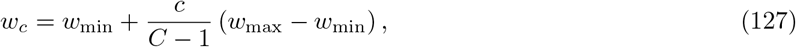

where *f*_*c*_, *P*_*c*_, and *w*_*c*_ denote the nominal stimulation frequency, number of pulses per burst, and pulse phase width assigned to class *c*, respectively. Where required by the stimulation hardware, these values were rounded or quantized to supported discrete settings.

The stimulation-current magnitude was held constant across classes during a given experiment. Each stimulation pulse consisted of a charge-balanced biphasic waveform,

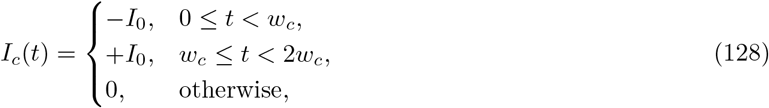

where *I*_0_ denotes the stimulation-current magnitude. The pulse waveform was repeated *P*_*c*_ times at frequency *fc*.

The complete input class was therefore represented by

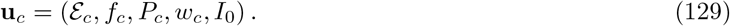

Thus, different classes perturbed the neuronal reservoir through distinct combinations of spatial stimulation location and temporal stimulation structure, while the stimulation-current magnitude was controlled independently.

### 4.7.2 State-conditioned presentation of input classes

Because the response of the neuronal reservoir depends on its instantaneous dynamical condition, input presentation was coupled to the state-awareness procedure described in Section 4.4. The instantaneous reservoir state was continuously estimated from recent spontaneous population activity, and a stimulation event was permitted only when the network occupied the target operating regime,

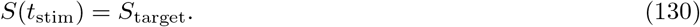

In the state-aware experiments considered here, the target regime was the operationally defined near-chaotic state. Once both the state criterion and the required inter-stimulation timing criterion were satisfied, the next input class was applied.

Input classes were presented repeatedly throughout the experiment, with the class index advanced after each accepted stimulation event. Consequently, the timing of individual trials was governed by the evolving biological state of the neuronal culture rather than by a completely fixed external schedule. The stimulation record retained the applied class identity together with its spatial and temporal stimulation parameters, enabling each measured neuronal response to be associated unambiguously with the corresponding input condition.

#### 4.7.3 Reservoir-state construction

For every accepted presentation *r* of class *c*, detected neuronal events were aligned to the corresponding stimulation time. Following the adaptive spike-detection procedure described in Section 4.3, spikes were integrated over a finite post-stimulation response window.

For recording electrode *i*, the event count associated with class *c*, trial *r*, threshold factor *k*, and response-window duration *W* was

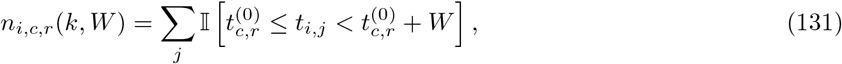

where 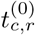 denotes the beginning of the response-analysis interval, *t i,j* is the time of detected event *j* on recording electrode *i*, and I [·] denotes the indicator function.

The trial-wise reservoir state was then represented by the population vector

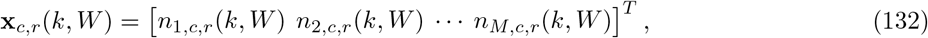

where *M* denotes the number of recording electrodes included in the separability analysis.

Recording electrodes used directly for stimulation could be excluded from the output population when required in order to reduce direct stimulation-site contamination. Denoting the candidate recording-electrode set by *C* and the stimulation-electrode set by *S*_stim_, the non-stimulating output population can be written as

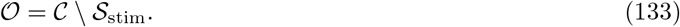

Each trial was associated with its known input label, producing the labelled reservoir-state dataset

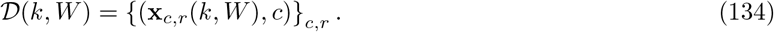

Importantly, the downstream separability analysis operated on the neuronal population response **x**_*c,r*_ rather than directly on the electrical stimulation parameters.

#### 4.7.4 Response-extraction parameter space

The experimentally observed geometry of the reservoir state depends partly on how neuronal events are extracted from the extracellular recordings. Separability was therefore evaluated over combinations of the adaptive spike-detection threshold factor *k* and the post-stimulation response-window duration *W* .

For every parameter pair,

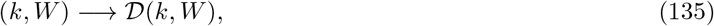

the corresponding set of trial-wise population states was reconstructed and evaluated independently. This generated a two-dimensional separability landscape over the response-extraction parameter space.

Evaluating a range of (*k, W* ) combinations serves two purposes. Variation in *k* tests whether class differentiation is preserved when the statistical stringency of spike detection is modified, whereas variation in *W* tests whether separability persists across different temporal integration scales of the evoked population response. Broad regions of high separability therefore indicate that the distinction among reservoir states is not restricted to a single narrowly tuned measurement condition.

#### 4.7.5 Class discrimination

Let *ĉ*_*r*_ denote the class predicted from the neuronal reservoir state of trial *r*, and let *c*_*r*_ denote its true stimulation class. Overall multiclass discrimination performance can be written as

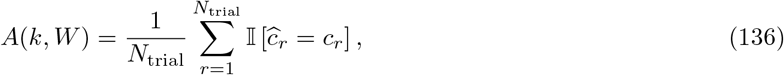

where *N*_trial_ is the number of evaluated reservoir responses.

Class-specific overlap was further quantified using the confusion matrix. For true class *a* and predicted class *b*,

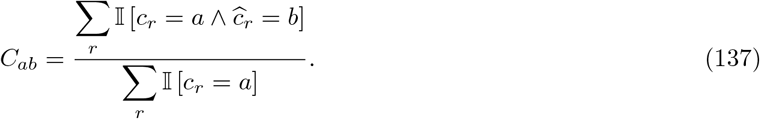

Diagonal entries therefore quantify class-specific correct discrimination, whereas off-diagonal entries quantify overlap between stimulation-induced reservoir representations.

For *C* equally represented input classes, the nominal random-classification level is

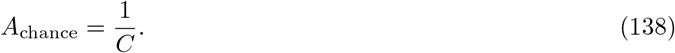

The measured discrimination can therefore be compared with the corresponding chance level when evaluating how effectively the reservoir transforms different inputs into distinguishable population states.

#### 4.7.6 Scaling of the input space

The same experimental construction was used while increasing the number of distinct spatio-parametric input classes. For each input-set size, stimulation locations and temporal stimulation configurations were assigned to the corresponding classes, and the resulting neuronal population responses were evaluated using the same reservoir-state construction and separability procedure.

Increasing *C* progressively increases the density of input representations that must be accommodated by the neuronal state space while simultaneously reducing the random-classification baseline,

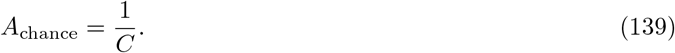

Comparison of the resulting separability landscapes and confusion structures therefore provides a direct assessment of whether the neuronal reservoir continues to map increasingly diverse spatio-parametric perturbations onto distinct regions of its accessible population-state space.

### 4.8 Fading-memory analysis

Fading memory was quantified from the relaxation of the multichannel neuronal state following the termination of each stimulation burst. Finite memory of recent inputs is a defining computational property of recurrent dynamical reservoirs [35] and has been observed experimentally in dissociated cortical networks and biological reservoir-computing systems [8, 9, 16]. The analysis was designed to measure how strongly an external perturbation displaced the reservoir from its local pre-stimulation state and how rapidly this displacement decayed after removal of the input. All temporal measurements were referenced to the end of the final stimulation pulse such that the subsequent dynamics represented the autonomous relaxation of the neuronal reservoir.

#### 4.8.1 Local baseline normalization

For each stimulation trial, a local pre-stimulation baseline was defined over the interval from −180 to −20 ms relative to stimulation onset. Because extracellular voltage amplitudes and background fluctuations vary substantially across MEA channels, each recording channel was normalized independently using robust baseline statistics, consistent with robust MAD-based normalization used for extracellular neural recordings [27, 29]. For channel *i*, the baseline median was calculated as

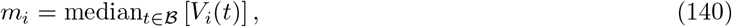

where *V*_*i*_(*t*) denotes the extracellular voltage recorded from channel *i* and ℬ denotes the pre-stimulation baseline interval. The corresponding robust estimate of the background scale was obtained from the median absolute deviation (MAD),

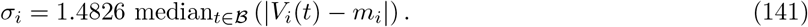

The voltage trace was subsequently expressed relative to its local background as

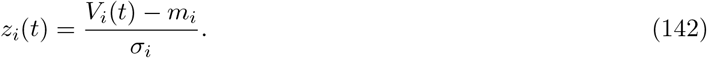

This channel-wise normalization places the activity recorded from different electrodes on a common dimensionless scale and reduces the influence of electrode-dependent differences in absolute extracellular voltage amplitude.

#### 4.8.2 Multichannel activity state

The post-stimulation response was divided into non-overlapping 20-ms windows. To reduce contamination from residual stimulation artifacts, the first 10 ms following the end of the stimulation burst was excluded. Analysis was continued to 230 ms after burst termination.

Within each temporal window centered at time *t*, the activity of channel *i* was represented by the root-mean-square (RMS) amplitude of its normalized voltage,

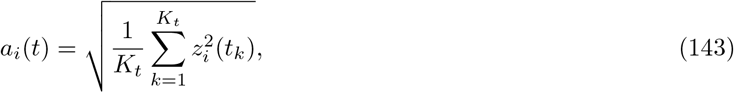

where *K*_*t*_ is the number of voltage samples contained within the corresponding 20-ms window. The baseline activity level of the same channel was similarly calculated over the pre-stimulation interval,

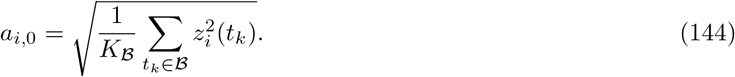

The stimulation-induced displacement of channel *i* from its baseline activity was therefore

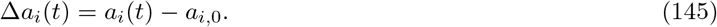

The instantaneous multichannel perturbation state was represented by the vector

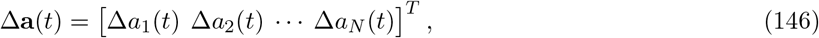

where *N* denotes the number of active recording channels.

#### 4.8.3 Population perturbation magnitude

The magnitude of the stimulation-induced perturbation was quantified as the RMS distance of the multichannel activity state from its local baseline state,

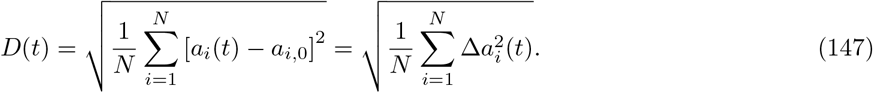

Thus, *D*(*t*) = 0 corresponds to a population activity envelope identical to the estimated baseline state, whereas increasing *D*(*t*) indicates a progressively larger population-level displacement from baseline. Because *D*(*t*) combines deviations across the electrode population, it provides a scalar measure of the magnitude of the evolving reservoir perturbation while retaining contributions from spatially distributed recording sites.

The initial perturbation magnitude, *A*_0_, was estimated as the mean perturbation magnitude over the first two valid post-blanking windows,

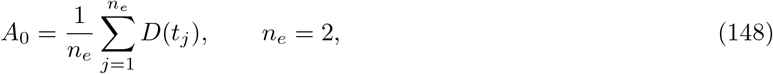

where *t*_*j*_ denotes the earliest measurable post-stimulation time points. Averaging the first two windows reduces sensitivity to fluctuations within a single temporal bin.

#### 4.8.4 Late-time residual and normalized fading-memory trace

The perturbation magnitude does not necessarily return exactly to zero within the finite observation period because of ongoing spontaneous activity and incomplete relaxation. A late-time residual level, *D*_floor_, was therefore estimated from the final two valid temporal bins,

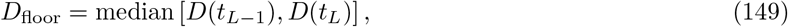

where *t*_*L*_ denotes the final analyzed time point.

The fading-memory magnitude was then normalized relative to the initial perturbation and late-time residual as

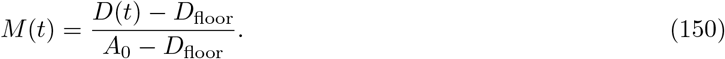

Under this normalization, *M* (*t*) ≈ 1 represents the early post-stimulation perturbation and *M* (*t*) ≈ 0 represents relaxation toward the late-time residual level. Values were constrained to the interval [0, 1.5] during construction of the empirical memory trace to limit the influence of transient excursions above the initial response. For fitting of the decay kernel, values were further constrained to [0, 1].

Time was shifted such that the first measurable post-blanking state corresponded to *t* = 0. Consequently, the fitted memory timescale describes relaxation from the earliest reliably measurable post-stimulation reservoir state rather than directly from the physical termination of the final stimulation pulse.

#### 4.8.5 Fading-memory timescale

The normalized memory trace was modeled using a stretched-exponential relaxation function,

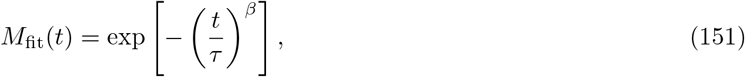

where *τ* is the characteristic relaxation timescale and *β* determines the shape of the forgetting kernel. For *β* = 1, Eq. 151 reduces to conventional exponential relaxation,

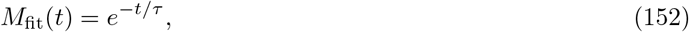

for which *M* (*τ* ) = *e*^*−*1^ ≈ 0.368. Thus, *τ* represents the time required for the fitted memory magnitude to decay to approximately 36.8% of its initial value when the relaxation is purely exponential.

Allowing *β* to vary accommodates departures from single-timescale exponential relaxation. Values *β <* 1 describe stretched decay with a comparatively long tail, consistent with relaxation distributed across multiple timescales, whereas *β >* 1 describes a more rapidly compressed decay. The parameters were estimated by nonlinear least-squares fitting with

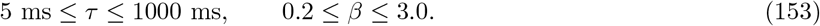

At least six valid post-stimulation temporal points were required for fitting.

#### 4.8.6 Memory persistence metrics

In addition to *τ*, memory persistence was quantified using empirical and model-derived threshold-crossing times. The empirical half-life, *t*_50_, was defined as the first interpolated time at which the measured normalized memory decreased to

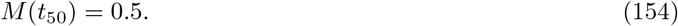

Similarly, *t*_20_ was defined by

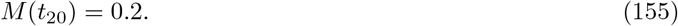

For the fitted stretched-exponential model, the analytical half-life follows from

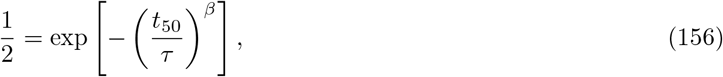

giving

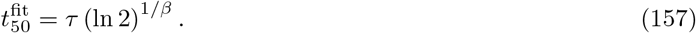

The integrated magnitude of memory over the observation interval was additionally quantified using the area under the normalized memory curve,

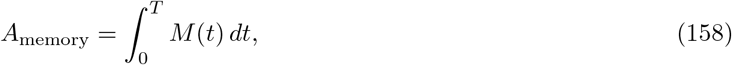

where *T* denotes the final analyzed delay. A temporal centroid was calculated to characterize where the memory mass was concentrated in time,

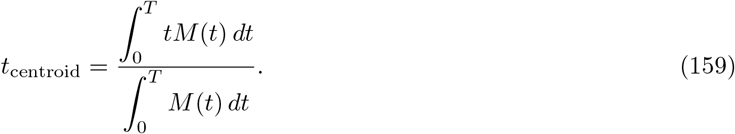

These complementary measures distinguish the characteristic decay timescale from the total magnitude and temporal distribution of the retained perturbation.

#### 4.8.7 Representational similarity to the early reservoir state

Because *D*(*t*) measures only the magnitude of displacement from baseline, an additional analysis quantified whether the *spatial pattern* of the population response remained similar to the initial post-stimulation configuration. The reference state was constructed by averaging the multichannel perturbation vectors from the first two valid post-stimulation windows,

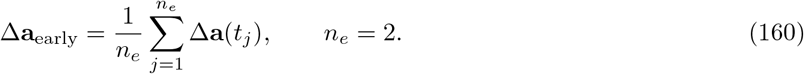

Representational persistence at subsequent time *t* was quantified using cosine similarity,

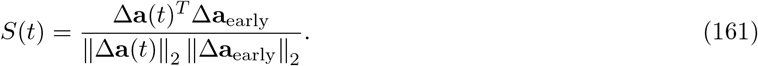

A value of *S*(*t*) = 1 indicates that the spatial pattern of the current perturbation is aligned with the early post-stimulation state, whereas decreasing similarity indicates progressive reorganization of the reservoir representation. This measure is conceptually distinct from *M* (*t*): *M* (*t*) quantifies how strongly the population remains displaced from baseline, whereas *S*(*t*) quantifies how closely the spatial structure of that displacement resembles the initial perturbation. The positive area under the similarity trace was additionally calculated as a measure of integrated representational persistence.

Together, *D*(*t*), *M* (*t*), *τ, t*_50_, and *S*(*t*) provide complementary descriptions of fading memory. *D*(*t*) measures the instantaneous magnitude of the stimulation-induced population displacement, *M* (*t*) expresses the normalized persistence of that displacement, *τ* and *t*_50_ quantify its characteristic lifetime, and *S*(*t*) measures preservation of the spatial population-state representation as the neuronal reservoir relaxes toward its ongoing spontaneous dynamics.

### 4.9 Mario Kart Control With a Neuronal Reservoir

Closed-loop control provides a direct test of whether living neuronal dynamics can support repeated perception– action computation, extending earlier demonstrations of cultured-neuron reservoir control and embodied biological computation [10, 13, 20].

#### 4.9.1 Game State and Reference Centerlines

The laptop client read the player position **p**_*i*_ = (*x*_*i*_, *z*_*i*_), checkpoint, continuous race-completion value, and elapsed client time from game memory. Each track was represented by an ordered centerline

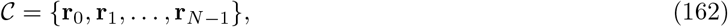

with segment lengths

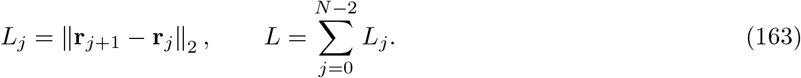

The client projected each observed position onto the nearest admissible centerline segment. A local search around the previous segment was used during normal motion; a global search was permitted when the local projection was more than 5000 game units from the kart. The projection returned the nearest point **q**_*i*_, segment index, cumulative arc position *s*_*i*_, and signed cross-track error *e*_*i*_. The normalized diagnostic error was

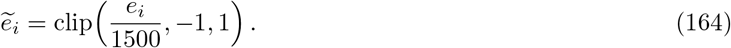

This error was logged but was not an input to the final signed-angle stimulation mapping.

The heading vector was calculated from consecutive kart positions when the estimated speed was at least 20 game units/s:

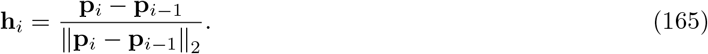

Before reliable motion was available, the tangent of the nearest centerline segment was used.

#### 4.9.2 Ghost-Point Heading Encoding

The normal ghost point was located 2600 game units ahead of the current arc position:

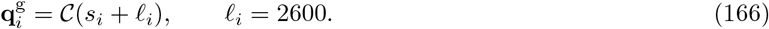

Let

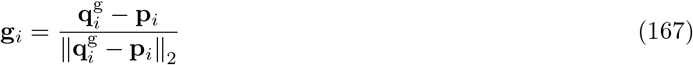

be the normalized direction from the kart to the ghost point. The client calculated

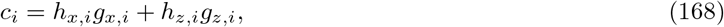

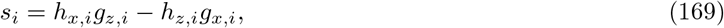

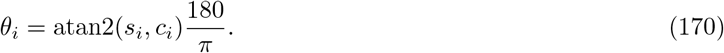

In the implemented local-coordinate form, 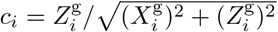 and 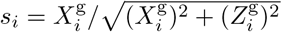. Positive *θ*_*i*_ indicates a rightward target and negative *θ*_*i*_ indicates a leftward target. The cosine and sine values were used only to recover this signed angle. They were not treated as separate neural inputs.

The lookahead changed only after a detected stuck state. The detector was not armed until the kart had first moved at least 300 game units/s. It then required speed at or below 120 game units/s continuously for 1.5 s. When this condition was satisfied, the target lookahead changed from 2600 to 1200 game units. The smoothed lookahead was updated as

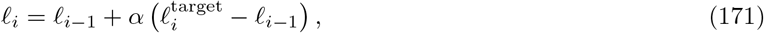

with *α* = 0.35 while shortening. Recovery required speed at or above 300 game units/s for 0.2 s and used *α* = 0.75 to restore the 2600-unit lookahead. The actual *ℓ*_*i*_ used for each request was logged.

#### 4.9.3 Mapping the Signed Angle to CL1 Stimulation

The angle was saturated at ±45° and normalized to

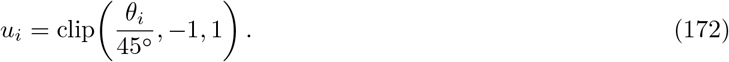

Stimulation frequency and current were

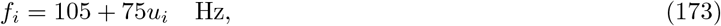

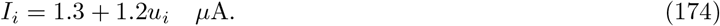

Therefore, *u* = −1 produced 30 Hz and 0.1 *µ*A, *u* = 0 produced 105 Hz and 1.3 *µ*A, and *u* = 1 produced 180 Hz and 2.5 *µ*A. Frequency and current changed together.

The chosen electrodes received the same stimulus in one stimulation call. There were no electrode-specific signs, offsets, phases, or secondary variables. Each stimulus contained three symmetric biphasic pulses. Each phase was 60 *µ*s, with a negative-current phase followed by an equal positive-current phase. The pulse repetition frequency was *f*_*i*_, and the magnitude of each phase was *I*_*i*_.

The expected five-class action was defined directly from the same signed angle:

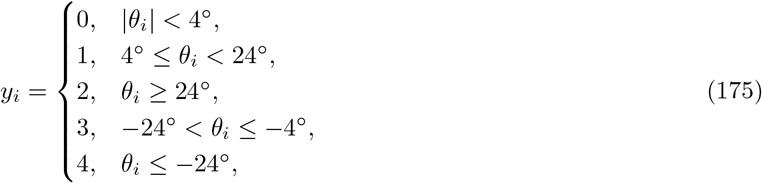

where classes 0–4 denote straight, soft right, hard right, soft left, and hard left. The expected class was logged for evaluation. It was not substituted for the live CL1 prediction.

#### 4.9.4 Readout Training and Model Selection

The final training collection contained 4000 CL1 responses, with 800 examples of each action class. Source states were sampled from the archived centerline controller data using random seed 42. When a class contained fewer than 800 unique states, source states were repeated to balance the plan. All responses derived from the same original state shared one group identifier so repeated states could not cross a training, validation, or test boundary.

A five-fold split assigned 3200 responses to model development and 800 responses to the held-out diagnostic set. Hyperparameters were selected within the development set by cross-validation, following established principles for avoiding circularity during neural-decoder evaluation and model selection [36]. The search evaluated the ten ridge penalties

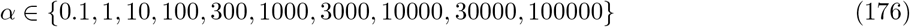

with either untransformed counts or log(1 + *x*) counts. Each pipeline applied median imputation, the selected count transform, feature standardization, and a class-balanced ridge classifier [37, 38]. Mean validation balanced accuracy [39] was the selection criterion. The selected fixed pipeline used log(1 + *x*) and *α* = 3000. The saved model also stored the ordered 60-column feature list, class map, encoding identifier, detector settings, source-file hash, and split provenance. The model was not updated during gameplay.

#### 4.9.5 Asynchronous Live Control

The laptop sampled the game at a nominal 5 Hz and enforced a minimum send interval of 0.200 s. Every UDP request contained a monotonically increasing request identifier, the signed-angle components, the expected class for later analysis, and the track/session identifiers. Up to eight unanswered requests could remain in flight. If several inference packets were already queued when the server became available, the server retained the newest game state and returned an explicit superseded-state error for older queued states. Requests older than 3 s were discarded by the client.

For an accepted request, the server validated the encoding identifier, the identical-eight-electrode mode, the detector parameters, and the 60 feature columns. It then stimulated CL1, extracted the response vector, evaluated the fixed ridge pipeline, and returned only the predicted action class. The client scheduled that returned class immediately. No additional 300-ms command delay was added. The expected class in the request was never used as a replacement for a missing or low-confidence prediction. The throttle key was held only after the first valid CL1 response was received.

Straight predictions released the steering key. Soft predictions generated a left or right steering pulse of

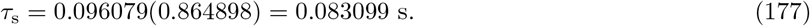

Hard predictions used

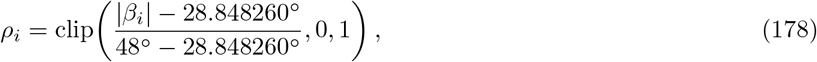

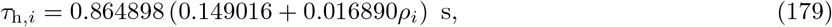

where *β*_*i*_ was the smoothed bearing to the longer controller target. Bearing smoothing used coefficient 0.687608. The minimum interval between executed commands was 0.013613 s. These steering values reproduce the optimized bot control profile. The CL1 round-trip supplied the communication and neural-response delay.

#### 4.9.6 Reset, Timeout, and Lap-Completion Rule

Each prospective attempt focused the game window, waited 0.5 s, pressed F1 for 0.12 s, and waited 3.0 s for the saved starting-line state to load. The completion detector used the game’s continuous `race_completion` field rather than centerline distance. The initial value, normally near 1.0 at the start of lap 1, defined the integer baseline *b*. A valid completion required sequential passage through relative progress milestones 0.20, 0.45, 0.70, and 0.90, followed by

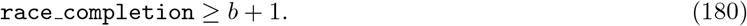

The final condition had to persist for three consecutive observations and could not end an attempt before 20 s. At 5 Hz, three confirmations add approximately 0.4 s after the first accepted finish sample. Centerline progress was retained for analysis but did not stop the prospective runs. Every attempt closed its server-side track session and flushed the per-track files before the next attempt.

#### 4.9.7 Centerline Tracking and Spatial Analysis

For consecutive observations, the time weight was

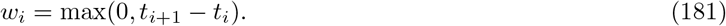

The final observation received zero weight because no subsequent interval was observed. The percentage of logged time inside the 1500-unit centerline band for run *r* was

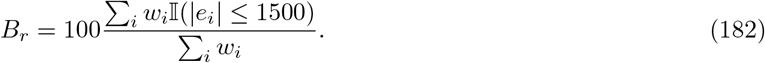

The 1500-unit threshold is a centerline-proximity.

For panels C and H, the centerline arc was divided into 40 equal-length sectors. Each observation was assigned to the sector of its nearest reference point. Within each run and sector *k*, the departure fraction was

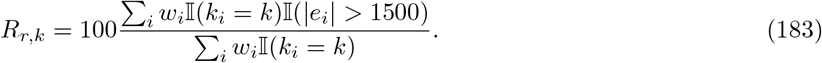

The displayed track map uses the unweighted mean of *R*_*r,k*_ over runs with an observed denominator in that sector.

For panel L, each multi-track observation was assigned by its logged fractional lap progress to one of 40 bins. The same time-weighted departure fraction was calculated separately for each track. Gray cells indicate that the trial did not contribute a valid observation to that sector. They are missing values, not zero tracking error.

Net centerline progress accumulated forward arc displacement with wrap-around. At crossing or nearby track sections, a proposed increment was rejected when it exceeded

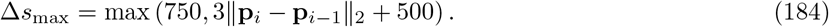

The uncapped final progress percentage was 100*S*_*r*_*/L*, where *S*_*r*_ is the accepted cumulative displacement. Panel K displays min(100, 100*S*_*r*_*/L*) so that travel beyond the nominal lap endpoint does not expand the vertical scale. The uncapped value remains in the derived data table.

#### 4.9.8 Lap-Time and Communication-Timing Definitions

The Track 1 game-clock values were recovered from the screen recording. For each of the ten sequential attempts, the first-lap split was the timer value held when the visible lap counter changed from (1/3) to (2/3). This boundary excludes screen-recorder lead-in, the period before the game timer started, the reset interval, and post-lap idling time. Panel E therefore reports video-verified in-game first-lap time. The video scan and final frame-level review preserved three decimal places from the displayed timer.

For request *i*, client round-trip time was

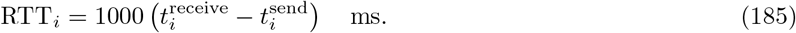

The run statistic was the median of its valid request-level RTT values. A cohort statistic was the median of the run medians, preventing a long run from receiving greater weight solely because it contained more requests. RTT spans client transmission, server queuing, stimulation and response acquisition, readout evaluation, return transmission, and client receipt. It is not an isolated biological latency.

## Acknowledgements

We thank Jack Gude for his part in developing the Mario Kart game interface to the CL1.

